# Modulation of SpyCatcher Ligation Kinetics by SpyTag Thioamide Substitution

**DOI:** 10.1101/2024.10.10.617615

**Authors:** Kristen E. Fiore, Denver Y. Francis, Sobrielle M. Casimir, Ryann M. Perez, E. James Petersson

## Abstract

Thioamide substitutions have been shown to impart valuable properties on peptides for biophysical experiments as well as cell or in vivo studies, but a rational understanding of thioamide effects on protein structure and protein-protein interactions is lacking. To elucidate their effects in β-sheet structures, we have used SpyCatcher003-SpyTag003 as a host-guest system to study individual thioamide incorporation at eight different positions in the SpyTag peptide. We have demonstrated that incorporating thioamides into SpyTag at specific positions can result in a ∼2-fold faster ligating complex, as well as >2000-fold slower ligating complex. Biophysical analysis and structural modeling provide a reasonable explanation for most of the thioamide effects, altering hydrogen bond networks as well as modulating an n→π* interaction within the SpyTag peptide. Our findings have important implications for potential applications of thioamide SpyTag variants, where the thioamide could impart protease stability in cells while also controlling the rate of ligation to SpyCatcher. These SpyCatcher-SpyTag host-guest experiments will also help to build a database for predicting thioamide effects on protein structure and function.

## Introduction

There are multiple isosteres of the peptide backbone that have been applied to make peptide mimics for biomedical applications, such as *N*-methyl amides, esters, (*E*)-alkenes, aza-amino acids, β-amino acids, triazoles and chloroalkenes.^1^ These isosteres can confer valuable properties on the peptide mimics through their differing hydrogen bond (H-bond) properties, conformational rigidity, and dipole moments. Of these isosteres, the thioamide is the most “sincere” mimic of the amide bond in that it maintains both a H-bond donor and acceptor without altering the stereochemistry of the α-carbon. Although only a single atom substitution, the O-to-S exchange has drastic effects on the chemical and physical properties of the peptide bond. The thioamide is larger due to the longer carbonyl bond and the increased van der Waals radius of sulfur. The lower oxidation potential allows thioamides to quench fluorophores in a PeT based mechanism.^2-4^ The red-shifted π-π* and n-π* absorptions result in unique circular dichroism (CD) signatures and the ability to quench UV fluorophores through a FRET mechanism.^5, 6^ This also results in a lower excitation energy for photoisomerization,^7^ so that the thioamide can be used as a *trans*-to-*cis* photoswitch.^8-11^

The thioamide is also a naturally occurring modification installed in various ribosomally synthesized and post-translationally modified peptides (RiPPs); such as in thioviridamide and methanobactin.^12^ To date, two proteins have been identified to contain thioamide post-translational modifications: methyl coenzyme M reductase (MCR)^13^ and the uL16 protein of the *E. coli* ribosome 70S subunit.^14^ Both proteins have been structurally characterized – MCR by X-ray crystallography and uL16 by cryo-electron microscopy The role of the thioamide in MCR is undetermined, although studies by Mitchell suggest that it increases the thermostability of the enzyme.^15^ Other proposals suggest that the thioamide could play a mechanistic role in the enzyme active site. For MCR, the thioamide is installed by a YcaO/TfuA enzymatic pair.^15, 16^ The thioamide in the uL16 protein was only recently discovered (2020) and has yet to be thoroughly investigated. Thus, although thioamides have been observed in structurally-characterized natural proteins, their role in those proteins is undetermined.

Due to the value of thioamides as biophysical tools and their enigmatic presence in natural proteins, our laboratory has undertaken a series of studies to better understand the effects of thioamides on protein secondary, teretiary, and quaternary structure. These investigations have involved the synthesis of peptides and proteins with thioamides at select locations, coupled with measurements of their stability and structure. Thioamides can be incorporated through solid-phase peptide synthesis (SPPS) as a nitro-benzotriazole activated *N*-Fmoc precursor.^17^ To incorporate a thioamide into a larger protein, an expressed or synthesized protein fragment can be ligated to the thioamide-containing peptide. Previously, our laboratory demonstrated this with native chemical ligation,^4, 18-20^ and we have optimized conditions for desulfurization of the Cys or Cys analog at the ligation site in the presence of a thioamide.^21^ This has allowed us to incorporate into proteins to quench fluorescence for the study of protein-peptide interactions,^4, 22^ protein folding,^4^ unfolding^23, 24^ or misfolding^18^. Our laboratory, as well as the Chatterjee laboratory, have collected thermodynamic data on the stabilities of thioamide-modified full-length proteins of various secondary structures.^25, 26^ We both found that seemingly similar positions had differing stabilities, particularly in β-sheet motifs. We collaborated to perform a systematic investigation of thioamide incorporation into two β-hairpin scaffolds complete with nuclear magnetic resonance (NMR) analysis.^27^ We found that the degree of perturbation was greatly dependent on the microenvironment of the residue. In particular, the right-handed twist of the β-hairpin allowed the thioamide to be perturbative as a H-bond acceptor at one position while neutral at another.

Empowered by these findings in model protein and peptide systems, we desired to explore thioamide effects in a wider variety of structural contexts. In our studies of thioamide inhibition of peptide cleavage, we were able to use a host-guest approach to create a database through combinations of thioamide peptide substrate guests and protease hosts.^28-30^ This database allowed us to develop a machine learning algorithm to predict the positional effects of thioamides in substrates of serine and cysteine protease with very high accuracy.^29^ We therefore considered protein systems that could be used to investigate thioamide effects in extended β-stands using a similar host-guest approach. One such system is SpyCatcher-SpyTag, a technology developed by the Howarth lab for protein labeling.^31^ Upon mixing the SpyCatcher protein fragment with the SpyTag peptide, an isopeptide bond forms that covalently links the two fragments together. Here, we synthesize SpyTag peptides with thioamides at a variety of positions and measure their rates of ligation with SpyCatcher. We find that some positions almost completely eliminate ligation, while others significantly increase the rate of ligation. We use additional biophysical and structural studies to gain initial understanding of these effects and set the stage for future host-guest studies with mutants of SpyCatcher as well as potential applications of thioamide-modified SpyTag peptides.

## RESULTS AND DISCUSSION

### Effects of Thioamidation on the Rate of SpyCatcher-SpyTag Complex Formation

To evaluate the impact of thioamidation on the association of an extended β-stand guest with a protein host, we utilized the SpyCatcher-SpyTag003 system, an improved variant of the original SpyCatcher-SpyTag001 system.^32^ This allowed us to systematically scan thioamide effects at sites which serve exclusive H-bond donor or acceptor roles within the core of the SpyCatcher binding pocket, as well as peripheral thioamide sites that would have more flexibility (**Fig. 1**). The peripheral thioamide sites included some that could potentially serve both H-bond roles and one that by definition can only serve as a H-bond acceptor, Val^S^_3_, which forms a tertiary thioamide with Pro_4_ (thioamide position denoted by a superscript “S”). We expected that core sites that served only as H-bond donors would stabilize complex formation, cores sites that served only as H-bond acceptors would destabilize complex formation, and peripheral sites would allow us to explore more nuanced effects of thioamide substitution.

**Figure 1.**
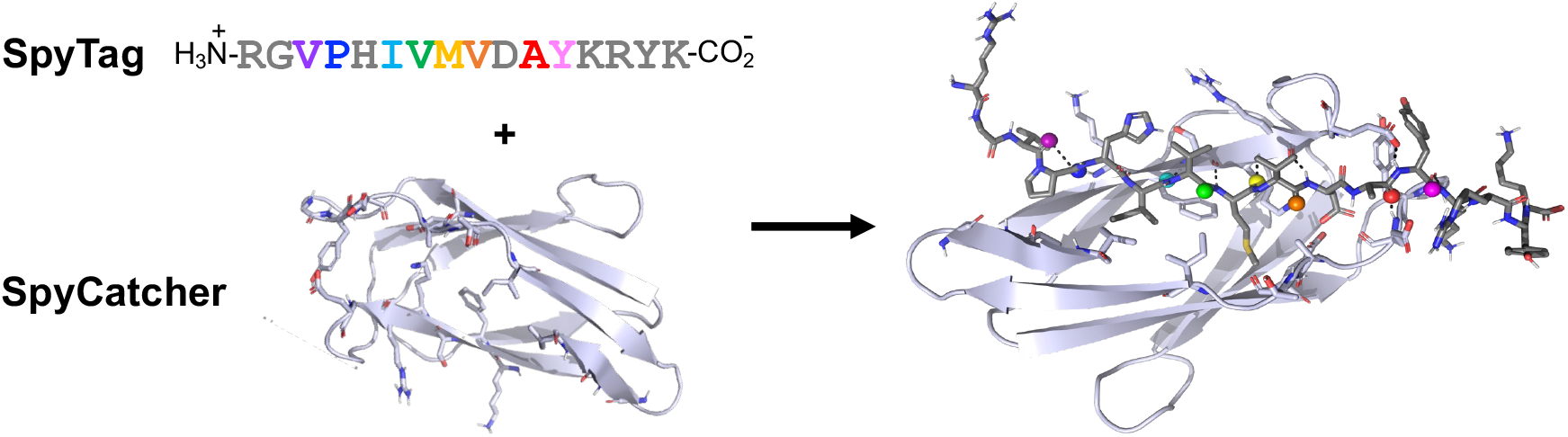
Ligation of Thioamide Variants of SpyTag Peptide with SpyCatcher Protein. Reaction scheme with positions of thioamide substitution in SpyTag colored, and overall structural model of SpyCatcher-SpyTag complex with thioamide positions highlighted, colored as in SpyTag sequence.

We synthesized these eight SpyTag003 variants, each with a single thioamide substitution, covering the majority of the SpyTag backbone. Peptides were synthesized on solid phase using established methods and purified by high performance liquid chromatography (HPLC) and the intact thioamide was verified by matrix-assisted laser desorption ionization mass spectrometry (MALDI MS). We also expressed SpyCatcher003 in *E. coli* and purified it using fast protein liquid chromatography (FPLC) for ligation reactions. The constructs will be simply referred to as SpyCatcher and SpyTag, unless explicit differences between the 001 and 003 systems are discussed.

Each SpyTag peptide was mixed with SpyCatcher protein for varying amounts of time, and then the reaction was quenched and product formation was evaluated using a gel shift mobility assay (**Fig. 2 Inset**, for all other primary gel data, see Supporting Information, SI). A progress curve was obtained from integration of the gel bands and fit to a second-order rate equation to obtain a rate constant (*k*_2_) for each thioamide variant as well as the all-amide SpyCatcher003 peptide, termed Oxo (**Fig. 2A** and **2B, Table 1**, see SI for individual curves, **Fig. S4**). From these data, we observed a wide variety of effects (**Fig. 2C**), from near-total inhibition of ligation (Met^S^_8_) to significant rate acceleration (Val^S^_3_ and Val^S^_7_).

**Table 1.**
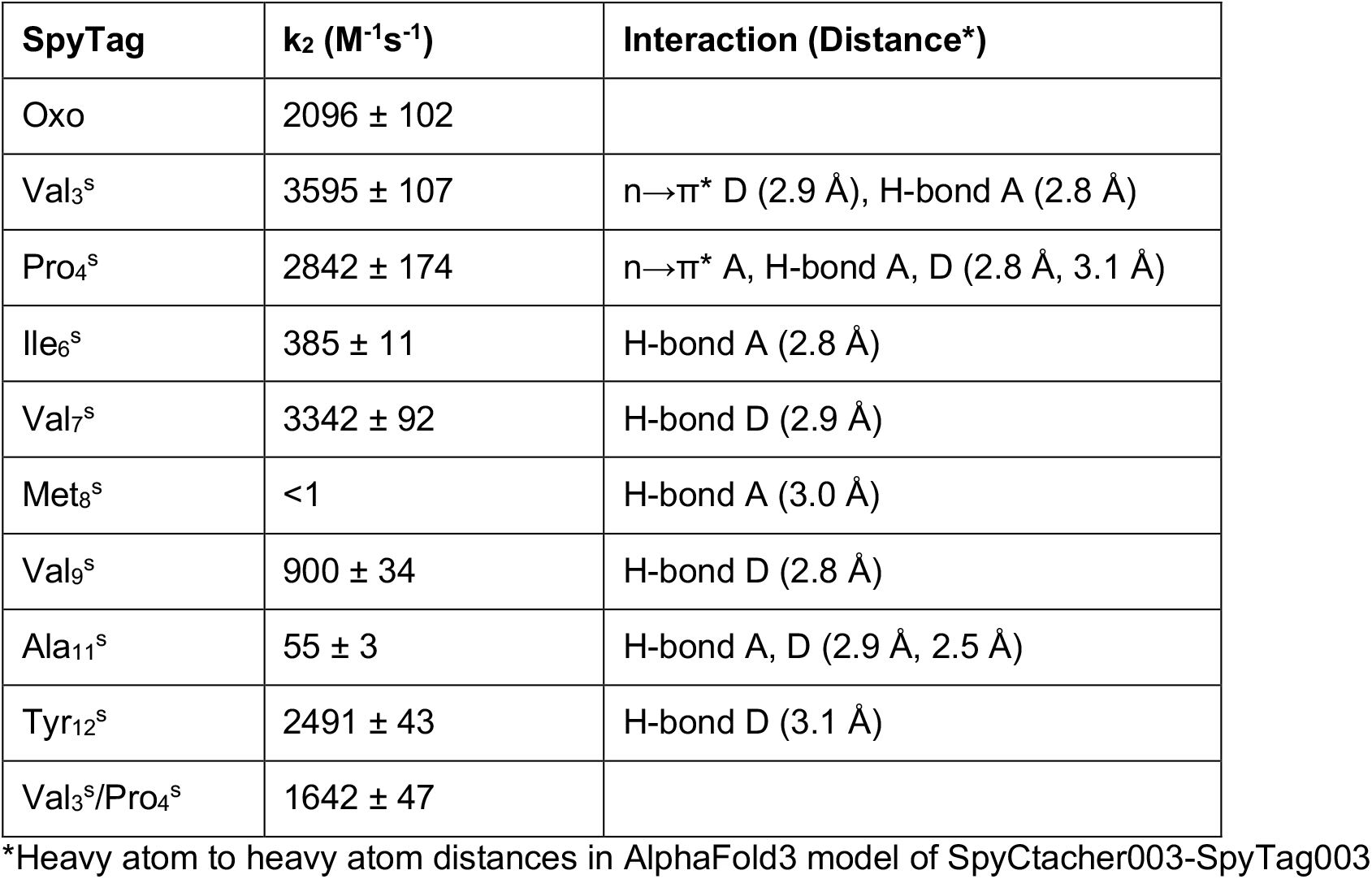
Gel Analysis Ligation Rates with Interaction Data from Structural Model.

| SpyTag | $k_2$ (M<sup>-1</sup>s<sup>-1</sup>) | Interaction (Distance*) |
| --- | --- | --- |
| Oxo | 2096 ± 102 |  |
| Val<sub>3</sub><sup>S</sup> | 3595 ± 107 | n→$\pi^*$ D (2.9 Å), H-bond A (2.8 Å) |
| Pro<sub>4</sub><sup>S</sup> | 2842 ± 174 | n→$\pi^*$ A, H-bond A, D (2.8 Å, 3.1 Å) |
| Ile<sub>6</sub><sup>S</sup> | 385 ± 11 | H-bond A (2.8 Å) |
| Val<sub>7</sub><sup>S</sup> | 3342 ± 92 | H-bond D (2.9 Å) |
| Met<sub>8</sub><sup>S</sup> | <1 | H-bond A (3.0 Å) |
| Val<sub>9</sub><sup>S</sup> | 900 ± 34 | H-bond D (2.8 Å) |
| Ala<sub>11</sub><sup>S</sup> | 55 ± 3 | H-bond A, D (2.9 Å, 2.5 Å) |
| Tyr<sub>12</sub><sup>S</sup> | 2491 ± 43 | H-bond D (3.1 Å) |
| Val<sub>3</sub><sup>S</sup>/Pro<sub>4</sub><sup>S</sup> | 1642 ± 47 |  |

\*Heavy atom to heavy atom distances in AlphaFold3 model of SpyCatcher003-SpyTag003

**Figure 2.**
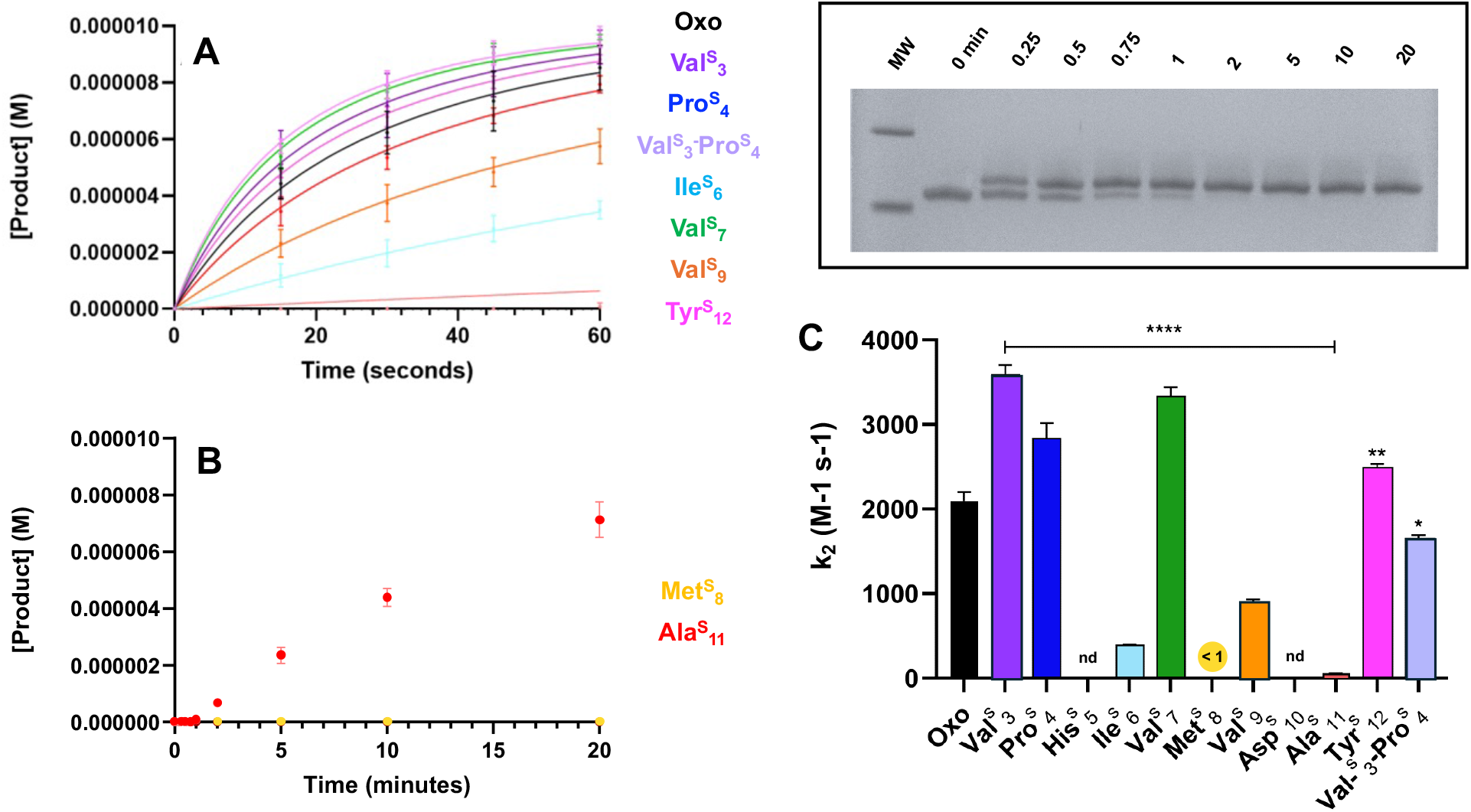
Thioamide SpyTag Ligation Rates. **A, B**. Progress curves for SpyCatcher and SpyTag association as determined by a gel shift assay. Assays were performed with 10 μM SpyCatcher003 and 20 μM SpyTag in 1X PBS, pH 7.5 at room temperature. **Inset**. Representative gel shift data for Oxo. **C**. Rate constants for thioamide SpyTag variants with statistical significance indicated. Significance was determined relative to Oxo using one-way ANOVA followed by Dunnett’s post hoc test; p < 0.05 (n = 8). Error bars represent the standard error of the mean (SEM) from triplicate assays. Points in panels A, B and bars in Panel C are colored according to Figure 1.

### Stopped Flow Analysis of SpyCatcher-SpyTag Constructs

To better understand the mechanistic basis for the thioamide effects, we turned to stopped flow fluorescence experiments, which could inform us as to whether each thioamide perturbs the initial rate of association or the chemical steps of ligation. Stopped flow experiments were enabled by the presence of a single Trp residue in SpyCatcher003, which could be quenched by thioamides that came within 20 Å upon complex formation.^21^ Based on our structural model of the SpyCatcher-SpyTag complex (see below), we anticipated that we would observe quenching of Trp fluorescence as the thioamide-containing SpyTag peptide associates with the Trp-containing SpyCatcher (**Fig. 3A** and **Table 2**). Val_3_, Ile_6_, Val_7_, and Val_9_ thioamide constructs were tested, spanning a range of ligation rates observed in gel ligation analysis. The Oxo peptide induced a change in fluorescence, allowing us to monitor its association, and all thioamide constructs generated significant quenching in comparison to Oxo (**Fig. 3B**). For all samples (including Oxo) except Ile^S^_6_, there is an initial rapid increase in fluorescence followed by a slower decrease. This two-step mechanism is consistent with previous biophysical studies of SpyCatcher-SpyTag.^33^ The initial increase is presumed to stem from changes in protein structure around Trp during preliminary complex association (Trp_58_ is in a hydrophobic core of Phe_30_, Phe_67_, Phe_76_ and Phe_93_, and is in proximity to Met_45_, which could change position to alter Trp quenching), with isopeptide bond formation occurring during the slower decay of fluorescence. Ile^s^_6_ differs in the initial phase, with a rapid decrease in fluorescence, followed by a slower second decreasing phase. The assignment of the second phase to the chemical step of ligation is supported by the similarity of the timescale for this fluorescence decrease to the gel data and the strong correlation between the relative rates for each thioamide construct when compared to the Oxo peptide (**Table 2**). In this light, the fluorescence increase observed for mixing of Ile^s^_6_ with SpyCatcher indicates that Ile^s^_6_ associates differently than the other peptides, which may set up the slower ligation process.

**Table 2.** Stopped-Flow Rate Data and Comparison to Relative Rates from Gel Analysis.

| SpyTag | 2<sup>nd</sup> Phase Rate (M<sup>-1</sup> s<sup>-1</sup>) | 2<sup>nd</sup> Phase Rate Constant (s<sup>-1</sup>) | Relative Rate Stopped Flow | Relative Rate Gel Analysis |
| --- | --- | --- | --- | --- |
| Oxo | 1.05 x 10<sup>-6</sup> | 0.042 ± 0.009 |  |  |
| Val<sup>S</sup><sub>3</sub> | 1.88 x 10<sup>-6</sup> | 0.075 ± 0.001 | 1.79 | 1.72 |
| Ile<sup>S</sup><sub>6</sub> | 2.00 x 10<sup>-7</sup> | 0.008 ± 0.005 | 0.19 | 0.18 |
| Val<sup>S</sup><sub>7</sub> | 1.45 x 10<sup>-6</sup> | 0.058 ± 0.018 | 1.38 | 1.59 |
| Val<sup>S</sup><sub>9</sub> | 6.50 x 10<sup>-7</sup> | 0.026 ± 0.012 | 0.62 | 0.42 |

**Figure 3.**
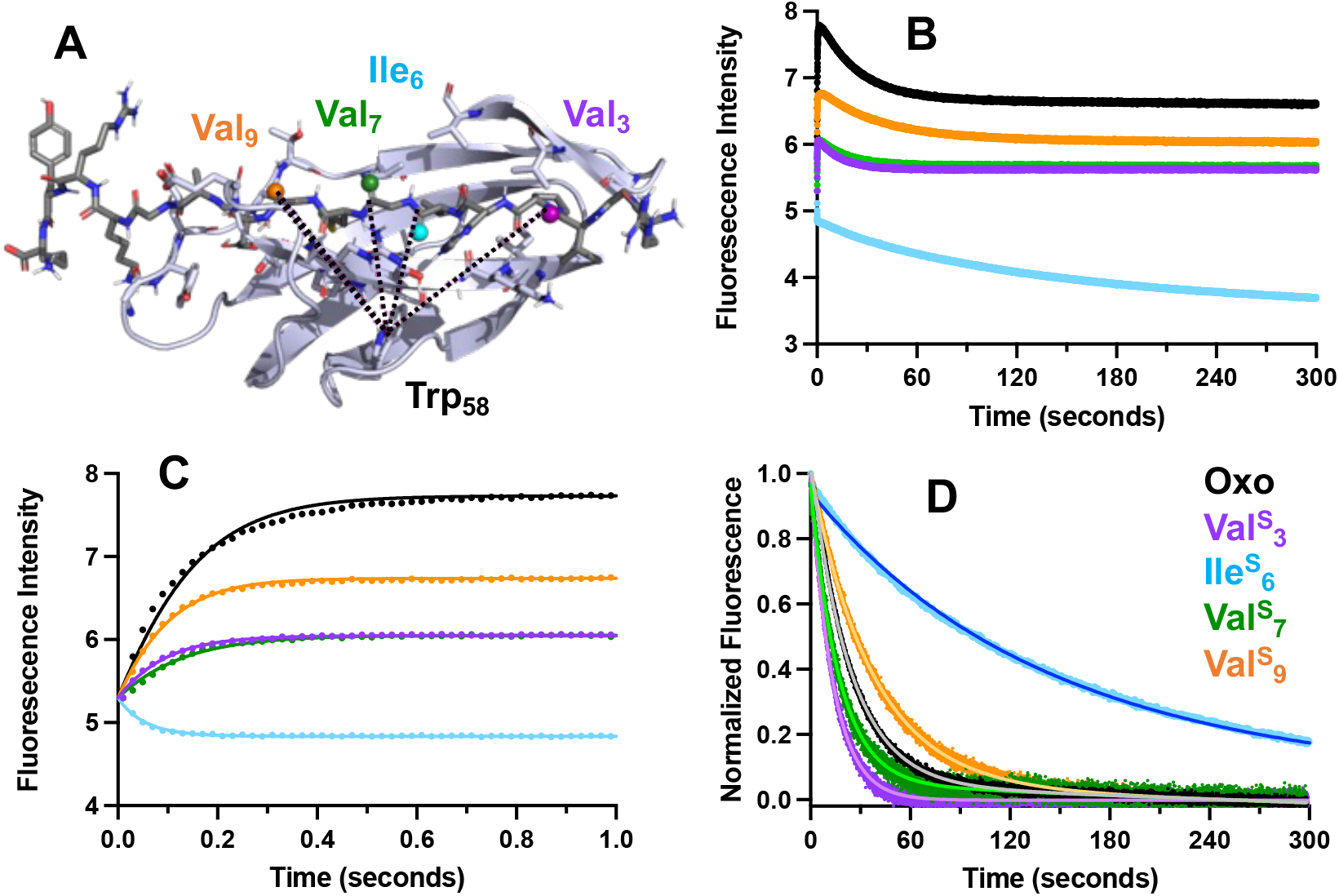
Trp fluorescence During SpyCatcher003 and SpyTag003 Complex Formation. **A**. Structural model showing positions of thioamide substitution relative to Trp_58_ fluorophore. **B**. Stopped-flow monitoring of Trp_58_ fluorescence changes during 25 μM SpyCatcher003 and 25 μM SpyTag peptide association in 1x PBS, pH 7.5 at room temperature (excitation at 295 nm and emission at 357 nm). **C**. Expanded view of first second of fluorescence trace showing changes due to initial binding. **D**. Normalized fluorescence for second phase fit to a first order decay to obtain rate constants reported in Table 2. All graphs colored according to legend in panel D.

### Biophysical Analysis of SpyCatcher-SpyTag Reaction

Before attempting to interpret the rate effects in terms of specific mechanisms such as changes in H-bond patterns, we wished to determine that the SpyTag variants were forming structurally similar protein complexes. A quick circular dichroism (CD) analysis of the ligation products for Oxo and five of the thioamide peptides showed no significant changes other than the presence of the thioamide n-to-π* absorbance at ∼270 nm (SI, **Fig. S5**). Intriguingly, the slower ligating Ile^S^_6_ has a larger n-to-π* CD band than the other mutants, perhaps indicating that it is in a more restricted electronic environment. However, Val^S^_3_ and Met^S^_8_ have comparable CD signatures with very different ligation kinetics, so there does not seem to be any clear correlation between the electronic environment of the thioamide and reactivity. The overall similarity of the CD data for all SpyCatcher-SpyTag complexes supported the pursuit of more detailed structural investigations.

To perform NMR studies of the complexes, we expressed and purified a ^15^N-labeled version of SpyCatcher, then mixed it with a SpyTag peptide and incubated with monitoring by MALDI MS to ensure that the isopeptide ligation reaction reached completion. The ligated SpyCatcher-SpyTag complex was then purified by FPLC and analyzed via ^1^H-^15^N heteronuclear single quantum coherence (HSQC) NMR experiments. We first compared the ^1^H-^15^N HSQC spectra of SpyCatcher alone and the Oxo SpyCatcher-SpyTag complex to verify that significant chemical shift changes could be observed (**Fig. 4**), consistent with previous NMR analysis of the SpyCatcher-SpyTag001 system.^33^ We then compared the spectra for the Oxo complex with the spectra for the Val^S^_3_, Val^S^_7_, and Met^S^_8_ complexes. We chose these three positions to include the two fastest thioamide variants and the slowest variant. Overlaying the ^1^H-^15^N HSQC spectra of the thioamide-containing complexes with Oxo complex spectra shows that there are no dramatic changes to the structures in the ligated complexes (**Fig. 4**). Met^S^_8_ spectra have some chemical shift perturbations compared to Oxo, but these are subtle. Thus, we concluded that analyzing the impacts of the thioamides using models based on the reported SpyCatcher-SpyTag structure seems to be justified.

**Figure 4.**
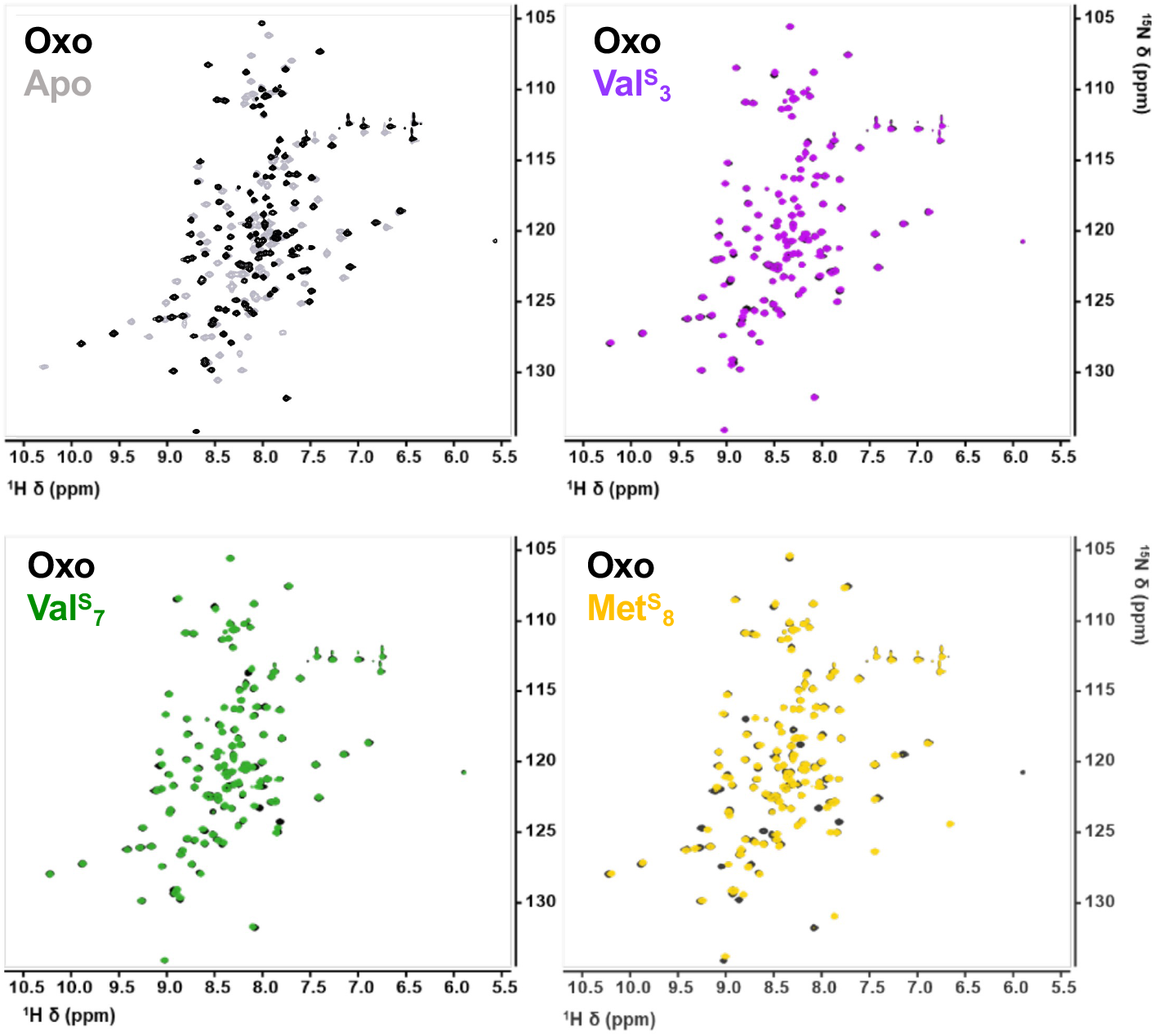
NMR Analysis of SpyCatcher-SpyTag. ^1^H-^15^N HSQC spectra of SpyCatcher only (Apo) or SpyCatcher-SpyTag complex for Oxo and thioamide variants. All three thioamide complexes display similar chemical shift patterns to Oxo despite significantly faster (Val^S^3, Val^S^_7_) or slower (MetS_8_) ligation rates.

### Structural Models of SpyCatcher-SpyTag Complex

We analyzed the crystal structure of the SpyCatcher001-SpyTag001 complex (PDB: 4MLI),^34^ as well as a computational model of the SpyCatcher003-SpyTag003 complex, for which no direct structural data is available. To validate modeling of SpyCatcher003-SpyTag003 using Alphafold3,^35^ we first constructed a model of SpyCatcher001-SpyTag001, and found that it aligned well with the published crystal structure (RMSD: 0.342 Å, SI, **Fig. S6**). The Alphafold2 prediction of the SpyCatcher003/SpyTag003 complex similarly aligned well with the SpyCatcher001-SpyTag001 structure (RMSD: 0.379). Hence, we do not believe that the presence of the thioamide in the various SpyTag003 constructs alters the chemical steps of ligation nor the folding of the SpyCatcher003/SpyTag003 complex. (**Fig. 5A**). The SpyTag001 peptide is identical to the SpyTag003 peptide for residues His_5_ to Tyr_12_, and there are no changes in the binding core between the two versions of SpyCatcher. Therefore, we have high confidence in our structure-based analysis for this region. An analysis of the local environment of each thioamide position shows that all of them except Val^S^_3_ have H-bond interactions with SpyCatcher with inter-heavy-atom distances of 2.5-3.2 Å (for both the experimental 001 structure and the 003 model), short enough that energetically significant H-bond interactions would be expected (**Fig. 5B, C, D** and **Table 1**). Thus, we will frame much of our interpretation of the ligation rate data in terms of H-bond effects.

**Figure 5.**
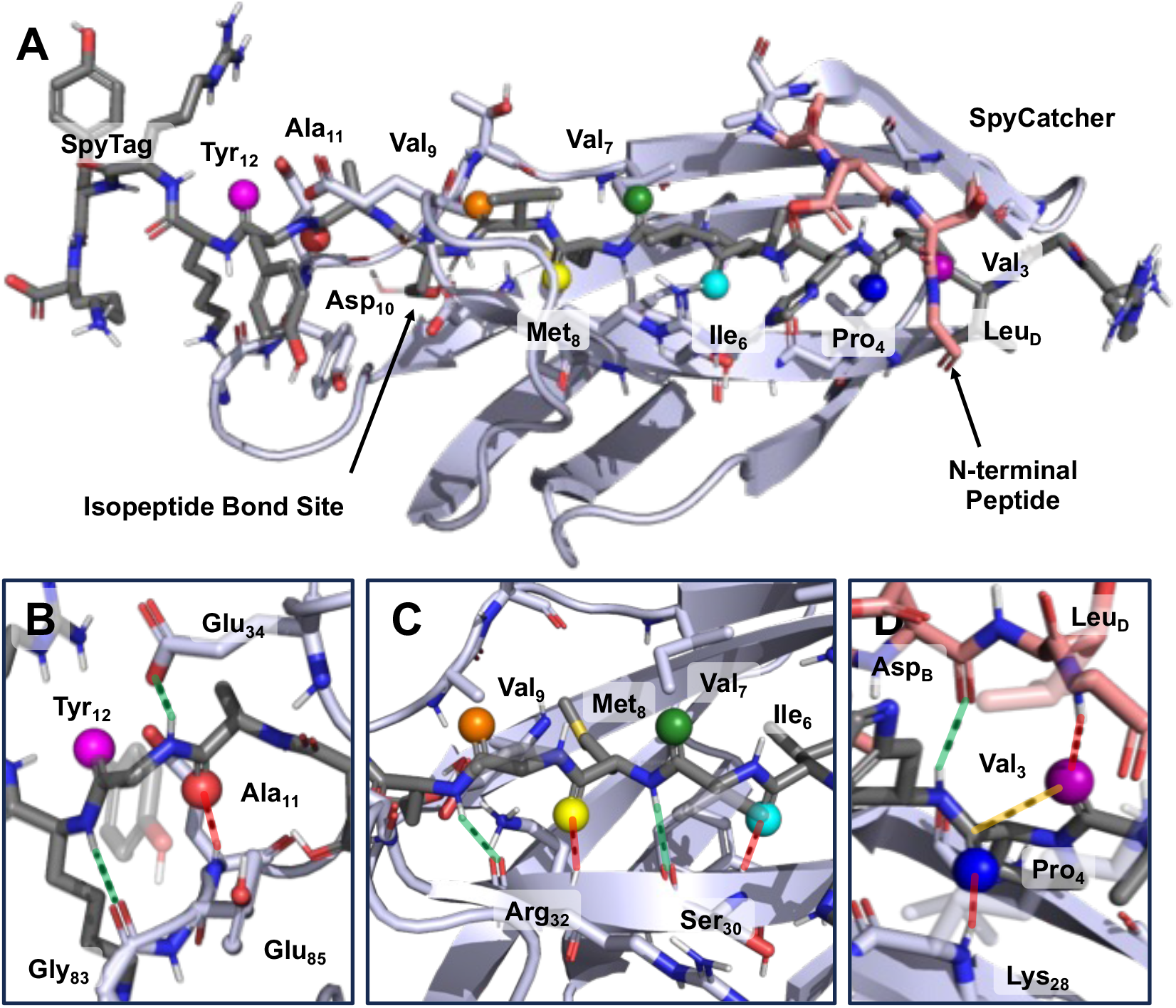
Structural Model of SpyCatcher003-SpyTag003 Complex. **A** Complete structure, constructed from Alphafold3 model of SpyCatcher003 protein (slate blue) with SpyTag003 peptide (grey) and CnaB2 domain N-terminal strand (salmon). Thioamide positions are colored to match ligation kinetics data in other figures with thiocarbonyl sulfur shown as a sphere. **B, C, D** Detailed structures highlighting H-bond interactions that could be disrupted (red dashed lines) or enhanced (green dashed lines) upon thioamidation, as well as an n→π* interaction (yellow).

Additional information comes from the crystal structure of the CnaB2 domain of FbaB, the fibronectin-binding protein of *Streptococcus pyogenes* from which the SpyCatcher-SpyTag system was derived.^31^ The CnaB2 domain structure (PDB ID: 2X5P)^36^ has a high degree of structural similarity to the SpyCatcher003-SpyTag003 Alphafold3 model (RMSD: 0.708 Å). However, one key difference is that while the N-terminus is unresolved in the SpyCatcher001-SpyTag001 structure and predicted with low confidence in the SpyCatcher003-SpyTag003 model, it can be seen clearly in the CnaB2 domain structure, wrapping around the substrate peptide (i.e. SpyTag), holding it in the pocket. Thus, by aligning the SpyCatcher003-SpyTag003 Alphafold3 model with the CnaB2 domain structure, we were able to add the N-terminal Val_A_-Asp_B_-Thr_C_-Leu_D_ peptide (**Fig. 5A**, salmon) into the SpyCatcher003 structure (**Fig. 5A**, slate blue) to identify additional H-bond interactions with Val_3_ and Pro_4_ (**Fig. 5D**).

### Mechanistic Analysis of Thioamide Interactions

When we designed the thioamide scanning experiment, we hypothesized that thioamides within the binding core positioned as exclusive H-bond donors (Val^S^_7_ and Val^S^_9_, **Fig. 5C**, green dashed lines) would accelerate association due to the more polarized N-H bond in the thioamide.^33^ Conversely, we expected core thioamides positioned as exclusive H-bond acceptors (Ile^S^_6_ and Met^S^_8_, **Fig. 5C**, red dashed lines) to slow association, as the sulfur of the thioamide is less electronegative and has additional steric bulk that might be perturbative. While our hypothesis that thioamides at H-bond acceptor positions would slow association proved correct, the difference in magnitude for the effects at Ile^S^_6_ and Met^S^_8_ (5.4-fold and >2000-fold, respectively) is quite surprising. The structures do not offer any obvious explanation for this difference as the H-bond distances are similar and neither site makes other significant contacts with the protein (**Fig. 5C** and **Table 1**). For Val^S^_7_ and Val^S^_9_, our H-bond donor principles did not even qualitatively explain the observed rates. While Val^S^_7_ enhanced ligation rates by 1.4-fold, similarly donating Val^S^_9_ ligated 1.6-fold slower than Oxo, potentially due to its adjacency to the isopeptide bond. In addition to affecting H-bond properties, thioamides are larger and have altered *cis*-*trans* kinetics and thermodynamics compared to amides, so these geometric factors may position Asp_10_ for more rapid isopeptide bond formation in the Val^S^_9_ complex. Consideration of the four peripheral thioamide positions can further our mechanistic understanding, since they all affected ligation rates significantly, even those that ostensibly made no interactions with the SpyCatcher protein.

Ala^S^_11_ ligates 38-fold slower than Oxo (**Table 1**), likely a consequence of being C-terminally adjacent to the isopeptide bond where stereoelectronic effects can inhibit catalysis similarly to Val^S^_9_. Ala^S^_11_ acts as both a H-bond donor and acceptor (**Fig. 5B**, green and red dashed lines), so it is likely that a local conformational change must occur to accommodate the larger thioamide in that H-bond sandwich, which may in turn change the active site geometry. Tyr^S^_12_ acts only as an H-bond donor and exhibits a mild rate acceleration (**Fig. 5B**, green dashed lines, and **Table 1**), consistent with our mechanistic model and its distance from Asp_10_. Thus, for thioamides at sites between Val_6_ and Tyr_12_, a general model emerges where H-bond donors accelerate ligation, H-bond acceptors retard ligation, and modifications in the vicinity of Asp_10_ uniformly disrupt ligation, presumably because of a high degree of sensitivity to structural changes in the catalytic site. In this context it is intriguing that Val^S^_3_ and Pro^S^_4_ modifications accelerate ligation. Both are far from the isopeptide bond and might be expected to have minimal impact. However, they are in a region that differs significantly between SpyTag003 and SpyTag001. The fact that replacement of the N-terminal Ala in SpyTag001 with a Val-Pro sequence (SI, **Fig. S6**) accelerates ligation supports the importance of this region and the potential for modulating ligation through thioamide substitutions there.

The rate acceleration for Val^S^_3_ (1.8-fold faster than Oxo, **Table 1**) is surprising in that it does not make any obvious interactions with the SpyCatcher protein in our Alphafold3 model and it cannot act as a H-bond donor since it is N-terminal to a Pro residue. However, the Val^S^_3_ construct potentially positions the thiocarbonyl to donate lone pair electrons to the Pro_4_ carbonyl in an n→π* interaction (**Fig. 5D**).^34, 35^ As demonstrated by Raines and coworkers, thioamides participate in stronger n→π* interactions than oxoamides.^36^ Since a stronger n→π* interaction stabilizes the *trans* form of that amide bond, we hypothesized that thioamide incorporation at the electron-donating Val^s^_3_ could lead to faster ligation by stabilizing a SpyTag peptide conformation that favors binding. Indeed, the Val-Pro substitution between SpyTag001 and SpyTag003 would be expected to favor such a conformation due to steric clash between the bulky sidechains and the thioamide n→π* effect would further accentuate it. Raines and colleagues also showed that an electron-accepting thioamide weakens the n→π* interaction due to decreased orbital overlap between thioamide acceptor and amide donor orbitals. Hence, if stabilizing the *trans* form of Val_3_-Pro_4_ accelerates ligation, one would expect that Pro^s^_4_ would ligate slower than Val^S^_3_, which it does (although still faster than Oxo, **Table 1**). Finally, since Raines demonstrated that incorporating thioamides into both the n→π* acceptor and donor positions enhances orbital overlap and further stabilizes the *trans* form, we hypothesized that double thioamide incorporation at Val_3_ and Pro_4_ would lead to even faster ligation. However, we found that the Val^S^_3_-Pro^S^_4_ dual thioamide peptide ligated 1.2-fold slower than Oxo and 2-fold slower than Val^S^_3_ (**Table 1**). Thus, these data suggest that if the n→π* interaction between Val_3_ and Pro_4_ is present, it is not the only factor governing ligation rate. For example, the Pro_4_ carbonyl may also accept an H-bond from the amide N-H of Lys_28_ in SpyCatcher, an interaction that is weakened upon thioamide substitution. An alternative explanation comes from examination of the crystal structure of the CnaB2 domain, and interactions with the flexible N-terminal tail.

The Val_A_-Asp_B_-Thr_C_-Leu_D_ segment of the CnaB2 domain N-terminal tail lays across the residues equivalent to Val^s^_3_-Pro^s^_4_ in the SpyTag peptide and makes H-bond contacts as well as less specific steric interactions (**Fig. 5D**). This four amino acid sequence and the intervening flexible linker are conserved in SpyCatcher003. Modeling this peptide fragment into the SpyCatcher003-SpyTag003 Alphafold3 model by aligning it with the CnaB2 domain structure shows the potential for SpyTag Pro_4_ amide N-H to donate a H-bond to the backbone carbonyl of SpyCatcher Asp_B_ and for SpyTag Val_3_ amide carbonyl to accept a H-bond from the backbone N-H of SpyCatcher Leu_D_. If we interpreted thioamide effects exclusively through these interactions, we would expect Pro^S^_4_ to be stabilizing and Val^S^_3_ to be destabilizing, which is not what we observed experimentally. However, if the effects from interactions with the Val_A_-Asp_B_-Thr_C_-Leu_D_ strand counterbalance the n→π* effects, then the moderate net stabilization effects of the monothioamides can be explained. The net destabilization by the dithioamide must be attributed to a disruption of the overall network of interactions by the increased steric bulk of the two adjacent thioamides.

Collectively, the structural models provide mechanistic explanations for the effects of the thioamides, where the extent of the effect is governed by the types of interactions and proximity to the isopeptide bond forming site, where sensitivity to a particular catalytic orientation can override the expected stabilizing effect of Val^S^_9_. Where thioamide sites are involved in multiple interactions, such as Ala^S^_11_ or the Val^S^_3_-Pro^S^_4_ dithioamide, destabilizing effects seem to dominate, perhaps due to steric incompatibility. A more complete understanding of the structural basis for these dominant destabilizing effects may be achievable through high resolution structural studies of the complexes through X-ray crystallography, mutagenesis of the SpyCatcher protein, and molecular dynamics simulations of SpyCatcher-SpyTag complex formation, all of which we are currently undertaking.

## Conclusion

We have utilized SpyCatcher003-SpyTag003 as a host-guest system to study the effects of individual thioamide incorporation at eight different positions in the SpyTag peptide. We have demonstrated that incorporating thioamides into SpyTag at specific positions can result in a ∼2-fold faster ligating complex (Val^S^_3_, Val^S^_7_), as well as >2000-fold slower ligating complex (Met^S^_8_). Low resolution structural characterization by CD and NMR showed the similarity of the SpyCatcher complexes with thioamide variants to the complex with Oxo, supporting an analysis of thioamide effects using structural models based on the existing crystal structure data. The AlphaFold3 model provides a reasonable explanation for most of the thioamide effects, particularly when one incorporates the N-terminal peptide which can be crystalographically observed in the CnaB2 domain structure. Previous biophysical studies have established that SpyCatcher-SpyTag association occurs through a two-step mechanism, but were unable to conclusively determine whether this conforms to an induced fit model or a conformational selection model, and imply that mutations can affect the mechanism to favor one model over another. This is consistent with our stopped-flow studies, where slower ligating Ile^S^_6_ appears to undergo a structurally-distinct initial association step based on its induced fluorescence increase. Intriguingly, the rate of the initial step is not that different, implying that the thioamide effect is more in setting up a geometry that leads to slow isopeptide bond formation. This has important implications for potential applications of thioamide SpyTag variants, where the thioamide could impart protease stability in cells while also controlling the rate of ligation to SpyCatcher. In addition to exploring such applications, we will use our established host-guest system to further understand thioamide effects on β-sheet systems by mutating the SpyCatcher protein in combination with thioamide modification. These experiments will help to build a database that can eventually be used to create a predictive model for thioamide effects on protein structure and function, a model applicable to both the design of synthetic thioamide peptides as well as understanding of the roles of thioamides in natural proteins.

## Acknowledgements

This work was supported by the University of Pennsylvania (Penn) and the National Science Foundation (NSF, CHE-2203909 to E.J.P.). K.E.F. and R.M.P. thank the National Institutes of Health (NIH) for funding through the Structural Biology & Molecular Biophysics Training Program (T32-GM-008275) and the Chemistry-Biology Interface Training Program (T32-GM-133398), respectively. D.Y.F thanks the University of Pennsylvania for funding through the Fontaine Society. The Bruker AVANCE NEO 600 MHz NMR spectrometer was supported by NIH supplement awards R01-GM-118510-03S1 and R01-GM-087605-06S1, and the Vagelos Institute for Energy Science and Technology. At Penn, the 600 MHz spectrometer is managed by Dr. Jun Gu and Dr. Chad Lawrence. K.E.F. acknowledges Dr. Chad Lawrence for his assistance with acquiring the NMR data at Penn and Mr. Ryan Kubanoff for assistance with the stopped-flow (supported by NSF, CHE-1337449). The matrix-assisted laser desorption ionization mass spectrometer was supported by NIH S10-OD-030460.

## References

1. Choudhary, A.; Raines, R. T., An Evaluation of Peptide-Bond Isosteres. ChemBioChem 2011, 12 (12), 1801–1807.

2. Bordwell, F. G.; Algrim, D. J.; Harrelson, J. A., The relative ease of removing a proton, a hydrogen atom, or an electron from carboxamides versus thiocarboxamides. J. Am. Chem. Soc. 1988, 110 (17), 5903–5904.

3. Petersson, E. J.; Goldberg, J. M.; Wissner, R. F., On the use of thioamides as fluorescence quenching probes for tracking protein folding and stability. Phys. Chem. Chem. Phys. 2014, 16 (15), 6827–6837.

4. Wissner, R. F.; Batjargal, S.; Fadzen, C. M.; Petersson, E. J., Labeling proteins with fluorophore/thioamide Förster resonant energy transfer pairs by combining unnatural amino acid mutagenesis and native chemical ligation. J. Am. Chem. Soc. 2013, 135 (17), 6529–6540.

5. Hollósi, M.; Kollát, E.; Kajtár, J.; Kajtár, M.; Fasman, G. D., Chiroptical labeling of folded polypeptide conformations: The thioamide probe. Biopolymers 1990, 30 (11-12), 1061–1072.

6. Sifferlen, T.; Rueping, M.; Gademann, K.; Jaun, B.; Seebach, D., β-Thiopeptides: Synthesis, NMR Solution Structure, CD Spectra, and Photochemistry. Helv. Chim. Acta 1999, 82 (12), 2067–2093.

7. Zhao, J.; Wildemann, D.; Jakob, M.; Vargas, C.; Schiene-Fischer, C., Direct photomodulation of peptide backbone conformations. Chem. Commun. 2003, (22), 2810–2811.

8. Helbing, J.; Bregy, H.; Bredenbeck, J.; Pfister, R.; Hamm, P.; Huber, R.; Wachtveitl, J.; De Vico, L.; Olivucci, M., A Fast Photoswitch for Minimally Perturbed Peptides: Investigation of the trans → cis Photoisomerization of N-Methylthioacetamide. J. Am. Chem. Soc. 2004, 126 (28), 8823–8834.

9. Satzger, H.; Root, C.; Gilch, P.; Zinth, W.; Wildemann, D.; Fischer, G., Photoswitchable Elements within a Peptide BackboneUltrafast Spectroscopy of Thioxylated Amides. J. Phys. Chem. B 2005, 109 (10), 4770–4775.

10. Bregy, H.; Heimgartner, H.; Helbing, J., A Time-resolved Spectroscopic Comparison of the Photoisomerization of Small β-Turn-forming Thioxopeptides. J. Phys. Chem. B 2009, 113 (6), 1756–1762.

11. Huang, Y.; Cong, Z.; Yang, L.; Dong, S., A photoswitchable thioxopeptide bond facilitates the conformation-activity correlation study of insect kinin. J. Pept. Sci. 2008, 14 (9), 1062–1068.

12. Mahanta, N.; Szantai-Kis, D. M.; Petersson, E. J.; Mitchell, D. A., Biosynthesis and Chemical Applications of Thioamides. ACS Chem. Biol. 2019, 14 (2), 142–163.

13. Grabarse, W.; Mahlert, F.; Shima, S.; Thauer, R. K.; Ermler, U., Comparison of three methyl-coenzyme M reductases from phylogenetically distant organisms: Unusual amino acid modification, conservation and adaptation. J. Mol. Biol. 2000, 303 (2), 329–344.

14. Watson, Z. L.; Ward, F. R.; Méheust, R.; Ad, O.; Schepartz, A.; Banfield, J. F.; Cate, J. H. D., Structure of the bacterial ribosome at 2 Å resolution. eLife 2020, 9.

15. Nayak, D. D.; Mahanta, N.; Mitchell, D. A.; Metcalf, W. W., Post-translational thioamidation of methyl-coenzyme M reductase, a key enzyme in methanogenic and methanotrophic archaea. eLife 2017, 6 (I), 1–18.

16. Mahanta, N.; Liu, A.; Dong, S.; Nair, S. K.; Mitchell, D. A., Enzymatic reconstitution of ribosomal peptide backbone thioamidation. Proc. Natl. Acad. Sci. U. S. A. 2018, 201722324–201722324.

17. Shalaby, M. A.; Grote, C. W.; Rapoport, H., Thiopeptide synthesis. α-Amino thionoacid derivatives of nitrobenzotriazole as thioacylating agents. J. Org. Chem. 1996, 61 (25), 9045–9048.

18. Batjargal, S.; Wang, Y. J.; Goldberg, J. M.; Wissner, R. F.; Petersson, E. J., Native chemical ligation of thioamide-containing peptides: Development and application to the synthesis of labeled α-synuclein for misfolding studies. J. Am. Chem. Soc. 2012, 134 (22), 9172–9182.

19. Wang, Y. J.; Szantai-Kis, D. M.; Petersson, E. J., Semi-synthesis of thioamide containing proteins. Org. Biomol. Chem. 2015, 13 (18), 5074–5081.

20. Fiore, K. E.; Phan, H. A. T.; Robkis, D. M.; Walters, C. R.; Petersson, E. J., Chapter Ten - Incorporating thioamides into proteins by native chemical ligation. In Methods Enzymol., Petersson, E. J., Ed. Academic Press: 2021; Vol. 656, pp 295–339.

21. Wang, Y. J.; Szantai-Kis, D. M.; Petersson, E. J., Chemoselective modifications for the traceless ligation of thioamide-containing peptides and proteins. Org. Biomol. Chem. 2016, 14, 6262–6269.

22. Goldberg, J. M.; Wissner, R. F.; Klein, A. M.; Petersson, E. J., Thioamide quenching of intrinsic protein fluorescence. Chem. Commun. 2012, 48 (10), 1550–1552.

23. Goldberg, J. M.; Batjargal, S.; Petersson, E. J., Thioamides as Fluorescence Quenching Probes: Minimalist Chromophores To Monitor Protein Dynamics. J. Am. Chem. Soc. 2010, 132 (42), 14718–14720.

24. Goldberg, J. M.; Speight, L. C.; Fegley, M. W.; Petersson, E. J., Minimalist probes for studying protein dynamics: Thioamide quenching of selectively excitable fluorescent amino acids. J. Am. Chem. Soc. 2012, 134 (14), 6088–6091.

25. Walters, C. R.; Szantai-Kis, D. M.; Zhang, Y.; Reinert, Z. E.; Horne, W. S.; Chenoweth, D. M.; Petersson, E. J., The effects of thioamide backbone substitution on protein stability: a study in α-helical, β-sheet, and polyproline II helical contexts. Chem. Sci. 2017, 8 (4), 2868–2877.

26. Khatri, B.; Majumder, P.; Nagesh, J.; Penmatsa, A.; Chatterjee, J., Increasing protein stability by engineering the n → π* interaction at the β-turn. Chem. Sci. 2020, 11 (35), 9480–9487.

27. Fiore, K. E.; Patist, M. J.; Giannakoulias, S.; Huang, C.-H.; Verma, H.; Khatri, B.; Cheng, R. P.; Chatterjee, J.; Petersson, E. J., Structural impact of thioamide incorporation into a β-hairpin. RSC Chem. Biol. 2022.

28. Barrett, T. M.; Chen, X. S.; Liu, C.; Giannakoulias, S.; Phan, H. A. T.; Wang, J.; Keenan, E. K.; Karpowicz, R. J., Jr.; Petersson, E. J., Studies of Thioamide Effects on Serine Protease Activity Enable Two-Site Stabilization of Cancer Imaging Peptides. ACS Chem Biol 2020, 15 (3), 774–779.

29. Giannakoulias, S.; Shringari, S. R.; Liu, C.; Phan, H. A. T.; Barrett, T. M.; Ferrie, J. J.; Petersson, E. J., Rosetta Machine Learning Models Accurately Classify Positional Effects of Thioamides on Proteolysis. J Phys Chem B 2020, 124 (37), 8032–8041.

30. Liu, C.; Barrett, T. M.; Chen, X.; Ferrie, J. J.; Petersson, E. J., Fluorescent Probes for Studying Thioamide Positional Effects on Proteolysis Reveal Insight into Resistance to Cysteine Proteases. ChemBioChem 2019, 20 (16), 2059–2062.

31. Zakeri, B.; Fierer, J. O.; Celik, E.; Chittock, E. C.; Schwarz-Linek, U.; Moy, V. T.; Howarth, M., Peptide tag forming a rapid covalent bond to a protein, through engineering a bacterial adhesin. Proc. Natl. Acad. Sci. U. S. A. 2012, 109 (12), E690–E697.

32. Keeble Anthony, H.; Turkki, P.; Stokes, S.; Khairil Anuar Irsyad, N. A.; Rahikainen, R.; Hytönen Vesa, P.; Howarth, M., Approaching infinite affinity through engineering of peptide– protein interaction. Proc. Natl. Acad. Sci. U. S. A. 2019, 116 (52), 26523–26533.

33. Zhang, N.; Liu, J.; Liu, Y.; Wu, W.-H.; Fang, J.; Da, X.-D.; Wang, S.; Zhang, W.-B., NMR Spectroscopic Studies Reveal the Critical Role of the Isopeptide Bond in Forming the Otherwise Unstable SpyTag–SpyCatcher Mutant Complexes. Biochemistry 2020, 59 (24), 2226–2236.

34. Li, L.; Fierer, J. O.; Rapoport, T. A.; Howarth, M., Structural Analysis and Optimization of the Covalent Association between SpyCatcher and a Peptide Tag. Journal of Molecular Biology 2014, 426 (2), 309–317.

35. Abramson, J.; Adler, J.; Dunger, J.; Evans, R.; Green, T.; Pritzel, A.; Ronneberger, O.; Willmore, L.; Ballard, A. J.; Bambrick, J.; Bodenstein, S. W.; Evans, D. A.; Hung, C.-C.; O’Neill, M.; Reiman, D.; Tunyasuvunakool, K.; Wu, Z.; Žemgulytė, A.; Arvaniti, E.; Beattie, C.; Bertolli, O.; Bridgland, A.; Cherepanov, A.; Congreve, M.; Cowen-Rivers, A. I.; Cowie, A.; Figurnov, M.; Fuchs, F. B.; Gladman, H.; Jain, R.; Khan, Y. A.; Low, C. M. R.; Perlin, K.; Potapenko, A.; Savy, P.; Singh, S.; Stecula, A.; Thillaisundaram, A.; Tong, C.; Yakneen, S.; Zhong, E. D.; Zielinski, M.; Ž㠄dek, A.; Bapst, V.; Kohli, P.; Jaderberg, M.; Hassabis, D.; Jumper, J. M., Accurate structure prediction of biomolecular interactions with AlphaFold 3. Nature 2024, 630 (8016), 493–500.

36. Oke, M.; Carter, L. G.; Johnson, K. A.; Liu, H.; McMahon, S. A.; Yan, X.; Kerou, M.; Weikart, N. D.; Kadi, N.; Sheikh, M. A.; Schmelz, S.; Dorward, M.; Zawadzki, M.; Cozens, C.; Falconer, H.; Powers, H.; Overton, I. M.; van Niekerk, C. A. J.; Peng, X.; Patel, P.; Garrett, R. A.; Prangishvili, D.; Botting, C. H.; Coote, P. J.; Dryden, D. T. F.; Barton, G. J.; Schwarz-Linek, U.; Challis, G. L.; Taylor, G. L.; White, M. F.; Naismith, J. H., The Scottish Structural Proteomics Facility: targets, methods and outputs. Journal of Structural and Functional Genomics 2010, 11 (2), 167–180.

