## Supporting Information for "Modulation of SpyCatcher Ligation Kinetics by SpyTag Thioamide Substitution"

### Materials and Methods

**General.** *Reagents.* *N*α-Fmoc-*N*ω-(2,2,4,6,7-pentamethyldihydro-benzofuran-5-sulfonyl)-L-arginine, 7-Azabenzotriazol-1-yloxy)tripyrrolidino-phosphonium hexafluorophosphate (PyAOP), triisopropylsilane (TIPS) were purchased from ChemImpex (Wood Dale, IL, USA). All other Fmoc-protected amino acids and resin were purchased from Novabiochem (currently Millipore Sigma; St. Louis, MO, USA). α-cyano-4-hydroxycinnamic acid (CHCA) was purchased from Santa Cruz Biotechnology, Inc (Dallas, TX, USA). *N*-methylmorpholine (NMM) and 1,8-Diazabicyclo[5.4.0]undec-7-ene (DBU) were purchased from Acros (currently Fisher Scientific; Waltham, MA, USA). All other reagents and solvents were purchased from Fisher Scientific or Millipore Sigma unless otherwise specified. Milli-Q filtered (18 MΩ) water was used for all solutions. All reagents and solvents were used without further purification. *Instrumentation.* RP-HPLC purification was performed on an Agilent 1260 Infinity II Preparative HPLC (Santa Clara, CA, USA). RP-HPLC analytical monitoring was performed on an Agilent 1260 Infinity II Analytical HPLC (Santa Clara, CA, USA). Matrix-assisted laser desorption/ionization time-of-flight (MALDI-TOF) mass spectra were collected with a Bruker MicroFlex (Billerica, MA, USA). NMR data were acquired with a Bruker AVANCE NEO 600 MHz spectrometer. Ultraviolet-visible (UV-vis) absorption spectra were collected on a GENESYS 150 UV-vis spectrophotometer (Thermo Fisher Scientific; Waltham, MA, USA). Circular dichroism (CD) data were acquired with a Jasco J-1500 CD spectrometer. Stopped-flow experiments were performed with a AutoSF120 stopped-flow spectrometer from KinTek (Snow Shoe, PA, USA).

#### 1. Synthesis and Characterization of Thioamide Precursors

*N*α-Fmoc-L-thiovaline-nitrobenzotriazolide was synthesized as previously reported.[41] *N*α-Fmoc-L-thioisoleucine-nitrobenzotriazolide[27] was synthesized as previously reported.[29]

*Synthesis and Purification of ThioMethionine (Fmoc-Met$^{S}$-Nbt) Precursor*

**Scheme S1. Fmoc-Met$^{S}$-Nbt (S1C) thioamide precursor synthesis**.

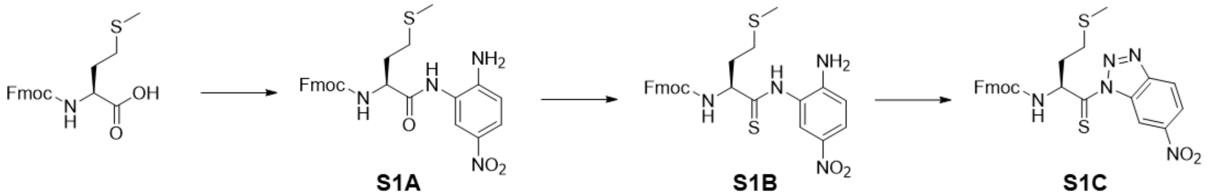


*N*α-Fmoc-L-thiomethionine-nitrobenzotriazolide was synthesized as previously described[42] with the following modifications.

*Coupling of the Fmoc-L Met with 1,2-diamino-4-nitrobenzene* (**S1A**). Fmoc-Met-OH (5 mmol, 1.86 g – 1 eq) was dissolved in dried THF (50 mL). *N*-methyl morpholine (10 mmol, 1.1 mL - 2 eq) was added and the reaction was cooled to -10 °C while purging with argon. Isobutyl chloroformate (IBCF, 5 mmol, 0.65 mL - 1 eq) was added dropwise to the stirred reaction. The syringe was rinsed with salt water from the bath to inactivate the residual IBCF. After 15 minutes stirring at -10 °C, 4-nitro-*o*-phenylenediamine (5 mmol, 0.766 g – 1 eq) was added to the reaction. The reaction was stirred under argon for 2 hours at -10 °C and then at room temperature overnight. After removing the solvent *in vacuo*, the product was dissolved in DMF (25 mL) and precipitated with the addition of 1:1 saturated KCl/MilliQ $H_2O$ (250 mL). The precipitate was filtered and washed extensively with water. The residual water was removed overnight under high vacuum.

*Preparation of the Fmoc L-amino ThioMet nitroanilides* (**S1B**). Phosphorous pentasulfide ($P_4S_{10}$, 2.25 mmol, 1.00g - 0.75 eq) and anhydrous $Na_2CO_3$ (2.25 mmol, 238.5 mg - 0.75 eq) were added to dry THF (30 mL) under argon and left stirring for 30 minutes (or until the phosphorous pentasulfide dissolved). After which, the compound **S1A** (3 mmol, 1.52 g – 1 eq) was added. The reaction was purged with argon and left to stir overnight. The next day, after removing the solvent *in vacuo*, the solid was resuspended in ethyl acetate and filtered over a pad of Celite©. The filtrate was washed twice with 5% $NaHCO_3$ and once with brine. The organic layers were combined and dried with $MgSO_4$. After filtration, the product was dissolved in minimal DCM and purified over silica on a Biotage Isolera One system (Biotage, LLC, Charlotte, NC, USA) with ethyl acetate/*n*-hexanes (25-75% ethyl acetate in n-hexanes).

*Preparation of Fmoc-L-amino ThioMet nitrobenzotriazolides* (**S1C**). The compound **S1B** (1.9 mmol, 1.004 g – 1 eq) was dissolved in 95% glacial acetic acid (25 mL) (v/v in Milli-Q water) and cooled to 0°C. $NaNO_2$ (2.9 mmol, 199 mg - 1.5 eq) was added slowly and the reaction was left to stir at 0 °C under atmosphere. After 30 minutes the product was precipitated with cold MilliQ $H_2O$. The filtered precipitate was lyophilized overnight and used directly for SPPS without further purification.

### 2. SpyTag Peptide Synthesis, Purification and Characterization.

***General.*** Peptides were manually synthesized via Fmoc SPPS on chlorotrityl resin (100-200 mesh, 1% DVB) from NovaBioChem in fritted syringes. All peptides were synthesized on either 50 or 100 µmol scale. At the end of the day, the resin was either stored dried or stirring in *N*-methyl-2-pyrrolidone (NMP).

The resin was swelled in *N*, *N*-dimethylformamide (DMF) or dichloromethane (DCM) (30-45 minutes). The first amino acid (Fmoc-Lys(Boc)-OH) (5 eq) was dissolved in DMF (2 mL or 4 mL) and *N, N*-diisopropylethylamine (DIPEA) (10 eq) was added. After vortexing, the mixture was added to the vessel. After stirring for 30 minutes at room temperature and washing with DMF (4 or 8 mL), the coupling was repeated. The unreacted resin was capped by treatment with DCM/MeOH/DIPEA (17:2:1 v/v) for 30 minutes. After draining and rinsing with DMF, the resin was deprotected with 20% v/v piperidine in DMF (2 or 4 mL) for 2 x 10 minutes. In-between all deprotections, the reaction vessel was washed with Wash 1 (4 or 8 mL each: DMF x2, DCM, DMF. After the last deprotection, the vessel the washed with Wash 2 (4 or 8 mL each: (DMF, DCM) x 3, DMF. The next amino acid (Fmoc-Tyr(tBu)-OH) (5 eq) and PyAOP (5 eq) were dissolved in DMF (2 mL or 4 mL) and DIPEA (10 eq) was added. After vortexing for at least 15 seconds, the mixture was added to the vessel. After stirring for 30 minutes at room temperature and washing with DMF (4 or 8 mL), the coupling was repeated. The deprotection was repeated as previously described. This series of couplings and deprotections continued until the thioamide was to be coupled.

***Thioamide Coupling and Deprotection.*** The nitrobenzotriazolide thioamide precursor (3 eq) was dissolved in anhydrous DCM stored over molecular sieves (purchased from Fisher) (1 - 2 mL). DIPEA (4 eq) was added, vortexed, and the mixture was added to the reaction vessel to stir at room temperature for 1 hour. After Wash 1, the coupling was repeated. The remaining unreacted termini were acetyl capped by treatment with 5 mL (8.4 mL DMF, 1.0 mL acetic anhydride, 0.6 mL NMM) for 2 x 10 minutes, with a DMF wash in-between. Following Wash 2, the Fmoc-group was removed with 2% DBU v/v in DMF (2 or 4 mL) for 3 x 2 minutes. After the first two DBU deprotections, Wash 1 was performed, and after the last deprotection, Wash 2 was performed.

The remaining amino acids were coupled as previously described; however, deprotections were performed with 2% DBU v/v in DMF (2 or 4 mL, 3x 5 minutes) to avoid thioamide residue epimerization and previously published side-reactions with piperidine.[43] Following removal of the Fmoc-group from $Arg_1$, Wash 2 was performed to clean the final product, followed by three washes with DCM (4 or 8 mL) and left under vacuum to dry the resulting resin.

***Cleavage.*** For $Val^S_3$, $Ile^S_6$, $Val^S_6$ and $Val^S_9$ peptides: a cleavage cocktail of 90% TFA, 2.5% 1,2-ethanedithiol (EDT), 2.5% TIPS, 2.5% thioanisole, 2.5% MilliQ $H_2O$ (v/v) (5 mL) was added to the vessel. For $Met^S_8$ peptide: a cleavage cocktail of 94% TFA, 2.5% EDT, 1 % TIPS, 2.5% MilliQ $H_2O$ (v/v) (5 mL) was added to the vessel. The vessel was rotated at room temperature for 45-60 minutes. The cleavage solution was drained from the syringe and rinsed with DCM or cleavage solution. After removal of the cleavage solution *in vacuo*, cold diethyl ether (20 mL) was added slowly to the product and the suspended precipitate was transferred to a 50 mL Falcon tube. The Falcon tube was cooled in dry ice for 15 minutes, and then the precipitate was collected by centrifugation at 4,000 RPM (3,313 x *g*) for 5 minutes. The diethyl ether was carefully poured off, and the diether ether wash (20 mL) was repeated. Following centrifugation and decanting, the pellet was left to dry in the hood overnight. Afterwards the pellet was stored at 4 °C.

***Purification.*** The crude peptide was dissolved in a minimal volume mixture of MilliQ $H_2O$ + 0.1% TFA, methanol and ≤ 20% acetonitrile (ACN) + 0.1% TFA and purified by RP-HPLC with a Luna Omega PS C18 preparative column (5 µm particle size, 250 mm length, 21.2 mm diameter) using the following gradients (**Table S1** and **Table S2**). The desired peptide was identified with MALDI MS using a CHCA matrix, and was dried on a lyophilizer (Labconco; Kansas City, MO, USA). The dried peptide was dissolved in MilliQ $H_2O$ + 0.1% TFA and subject to second-pass RP-HPLC using one of two columns: Luna Omega PS C18 preparative column (5 µm particle size, 250 mm length, 21.2 mm diameter) or a Phenomenex Luna Omega PS C18 semi-preparative column (5 µm particle size, 250 mm length, 10 mm diameter) using the following gradients (**Table S1** and **Table S2**). Purity of the peptides was assessed by analytical RP-HPLC with a Phenomenex Luna Omega PS C18 column (5 µm particle size, 150 mm length, 4.6 mm diameter) with gradient **H**. After purification SpyTag peptides were ≥ 90% pure (based on analytical 215 nm AUC integration).

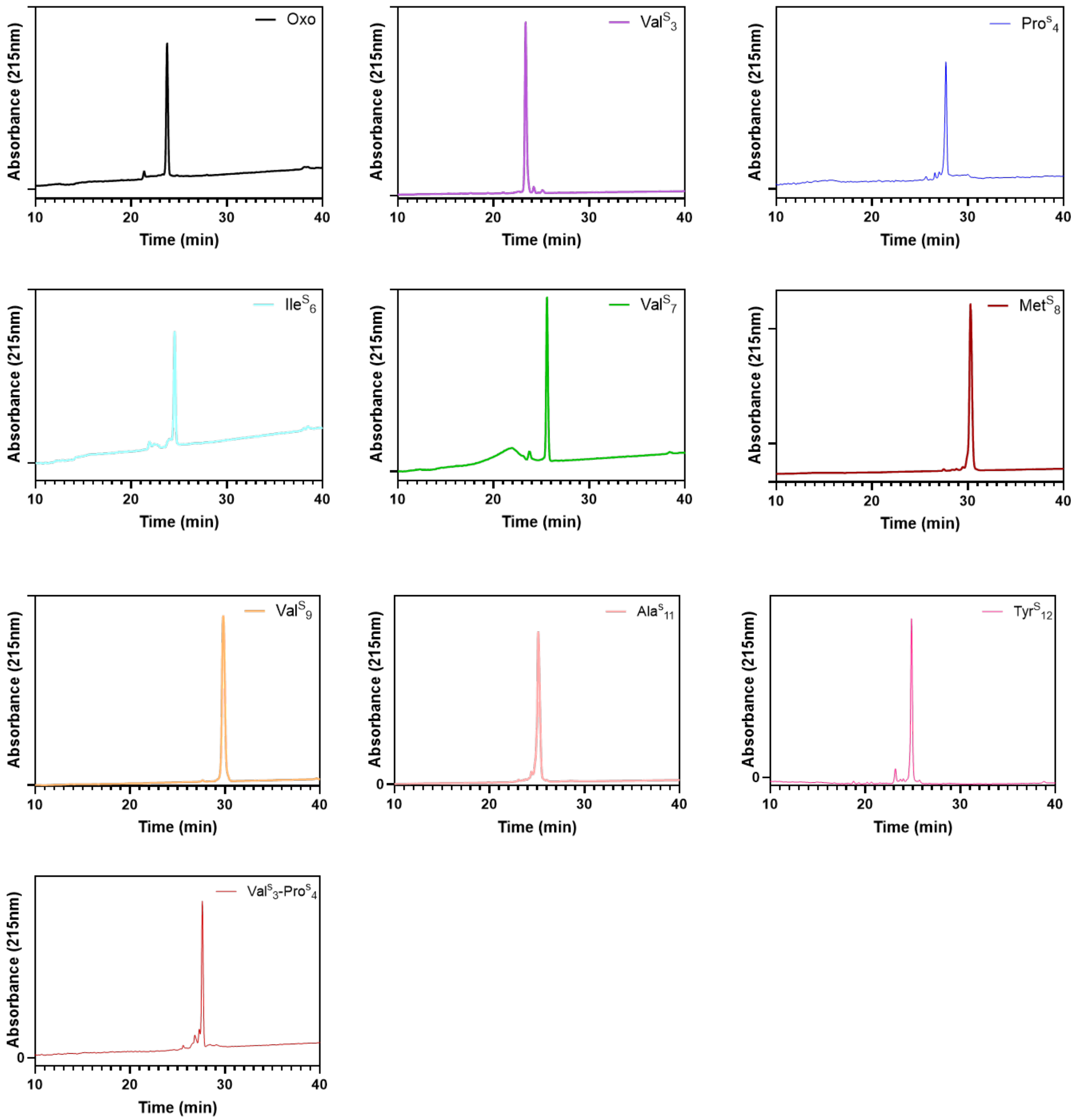


**Figure S1.** Analytical HPLC traces of synthesized SpyTag variants (gradient H).

**Table S1.** MALDI characterization, gradient of purification, and analytical retention time of the synthesized SpyTag peptides

| SpyTag Peptide | $[M+H]^+$ (m/z) | | $[M+Na]^+$ (m/z) | | $[M+K]^+$ (m/z) | | Gradient | Analytical Retention Time* |
|---|---|---|---|---|---|---|---|---|
| | Exp | Obs | Exp | Obs | Exp | Obs | | |
| WT | 1932.06 | 1931.95 | 1954.05 | 1953.96 | 1970.02 | 1969.96 | **A**, **B** | 23.75 |
| $Val^{S}_{3}$ | 1948.04 | 1948.06 | 1970.02 | 1970.03 | 1986.00 | 1986.02 | **A**, **C** | 23.35 |
| $Pro^{S}_{4}$ | 1948.04 | 1948.01 | 1970.02 | --- | 1986.00 | ----- | **A,B** | 27.71 |
| $Val^{S}_{3}$-$Pro^{S}_{4}$ | 1965.45 | 1964.94 | 1988.44 | --- | 2004.55 | ----- | **A,B** | 27.64 |
| $Ile^{S}_{6}$ | 1948.04 | 1947.88 | 1970.02 | 1969.86 | 1986.00 | ----- | **D**, **E** | 24.54 |
| $Val^{S}_{7}$ | 1948.04 | 1947.96 | 1970.02 | 1969.92 | 1986.00 | ----- | **A**, **B** | 25.59 |
| $Met^{S}_{8}$ | 1948.04 | 1948.75 | 1970.02 | --- | 1986.00 | ----- | **A**, **F** | 30.27 |
| $Val^{S}_{9}$ | 1948.04 | 1948.45 | 1970.02 | --- | 1986.00 | ----- | **A**, **G** | 29.32 |
| $Ala^{S}_{11}$ | 1948.04 | 1949.11 | 1979.02 | --- | 1986.00 | ----- | **H** | 25.13 |
| $Tyr^{S}_{12}$ | 1948.04 | 1948.03 | 1979.02 | --- | 1986.00 | ----- | **H** | 24.86 |

**Table S2.** HPLC gradients for purification (solvent A = MilliQ water with 0.1% TFA, solvent B = acetonitrile with 0.1% TFA)

| **Grad.** | **Time** | **% B** | **Grad.** | **Time** | **%B** |
|---|---|---|---|---|---|
| **A** | 0.00 | 5 | **B** | 0.00 | 5 |
| | 3.00 | 5 | | 3.00 | 5 |
| | 8.00 | 20 | | 8.00 | 18 |
| | 11.00 | 20 | | 11.00 | 18 |
| | 31.00 | 40 | | 33.00 | 40 |
| | 34.00 | 40 | | 36.00 | 40 |
| | 38.00 | 100 | | 40.00 | 100 |
| | 43.00 | 100 | | 45.00 | 100 |
| | 45.00 | 5 | | 48.00 | 5 |
| **Grad.** | **Time** | **% B** | **Grad.** | **Time** | **%B** |
| **C** | 0.00 | 5 | **D** | 0.00 | 5 |
| | 3.00 | 5 | | 3.00 | 5 |
| | 8.00 | 20 | | 8.00 | 20 |
| | 11.00 | 25 | | 11.00 | 20 |
| | 31.00 | 30 | | 31.00 | 35 |
| | 36.00 | 100 | | 34.00 | 35 |
| | 42.00 | 100 | | 38.00 | 100 |
| | 45.00 | 5 | | 43.00 | 100 |
| | | | | 45.00 | 5 |

| **Grad.** | **Time** | **% B** | **Grad.** | **Time** | **%B** |
|---|---|---|---|---|---|
| **E** | 0.00 | 5 | **F** | 0.00 | 5 |
| | 3.00 | 5 | | 3.00 | 5 |
| | 8.00 | 20 | | 8.00 | 25 |
| | 11.00 | 20 | | 11.00 | 25 |
| | 31.00 | 30 | | 31.00 | 35 |
| | 33.00 | 30 | | 33.00 | 35 |
| | 37.00 | 100 | | 37.00 | 100 |
| | 42.00 | 100 | | 45.00 | 100 |
| | 45.00 | 5 | | 48.00 | 5 |
| **Grad.** | **Time** | **% B** | **Grad.** | **Time** | **%B** |
| **G** | 0.00 | 5 | **H** | 0.00 | 5 |
| | 3.00 | 5 | | 3.00 | 5 |
| | 8.00 | 20 | | 8.00 | 12 |
| | 11.00 | 20 | | 11.00 | 12 |
| | 31.00 | 25 | | 51.00 | 35 |
| | 33.00 | 30 | | 54.00 | 35 |
| | 37.00 | 100 | | 58.00 | 100 |
| | 42.00 | 100 | | 63.00 | 100 |
| | 45.00 | 5 | | 65.00 | 5 |

### 3. SpyCatcher003 Cloning, Expression and Purification

#### *1.3.1 Cloning*

***General.*** Cloning was performed in a T100 thermocycler from Bio-Rad (Hercules, CA, USA). Primers and double stranded DNA (gblocks) were ordered from Integrated DNA technologies (Coralville, IA, USA). Q5 Hot Start High Fidelity 2x Master Mix, 2x HiFi Assembly Master Mix, T4 DNA Polynucleotide Kinase, T4 DNA ligase and DH5α High Efficiency *E.coli* cells were from New England Biolabs (Ipswich, MA, USA). QIAprep Spin Plasmid Miniprep Kit and DNA Concentrator & Cleanup Kit were from Qiagen (Germantown, MD, USA). DNA concentrations were determined on a TECAN Nanoquant plate on a TECAN Infinite® M1000Pro plate reader (Männedorf, Switzerland). DNA sequencing was performed at the University of Pennsylvania Sequencing Core (Philadelphia, PA, USA).

***SpyCatcher003 Gibson Assembly***

The SpyCatcher003 sequence was codon optimized using the online GeneArt tool from ThermoFisher (Waltham, MA, USA). Primers were designed with 15+ nt overhangs that correspond to insert sequence (SpyCatcher003 gblock). The gblock was dissolved in sterile TE buffer to a concentration of 10 ng/μL. The primers were dissolved in DNAse/RNAse free water to a concentration of 50 μM. The parent plasmid (pTXB1 GB1-NpuDnaE-His$_6$) was PCR amplified with the overhang-containing primers with the following conditions (**Table S3**, annealing temperature of 61 °C, extension total time of 5 minutes, Q5 Hot Start High Fidelity 2x Master Mix (25.0 μL), forward primer (0.5 μL), reverse primer (0.5 μL), plasmid (diluted with DNAse/RNAse free water to < 1ng/μL – 1.0 μL)).

**Table S3.** Thermocycler PCR Settings

| Step Number | Temperature | Duration |
|---|---|---|
| 1 | 98 °C | 30 sec |
| 2 | 98 °C | 10 sec |
| 3 | __°C (Annealing temperature) | 30 sec |
| 4 | 72 °C (Extension) | 45 sec/ kb |
| 5 | Go to step 2 | 29 times |
| 6 | 72 °C | 2 min |
| 7 | 4 °C | Hold |

Following PCR amplification, the PCR reaction was cleaned-up with a DNA Concentrator & Clean-up Kit and eluted with 10 μL. The resulting DNA was quantified. The PCR amplified plasmid (vector) was assembled with the gblock insert (100 ng vector, insert (5 eq), DNAse/RNAse free water to 10 μL and 2x HiFi Assembly Master Mix (10 μL)) through incubation at 50 °C for 60 minutes. The reaction was cooled on ice and 2 μL was added to an aliquot of DH5α High Efficiency cells (NEB, C2987). The assembled plasmid was transformed following NEB protocol. Individual colonies were isolated and grown in 5 mL LB medium overnight at 37 °C. The DNA was extracted using the Plasmid Miniprep Kit, quantified, and submitted for sequencing (T7 promoter).

**Gblock:**

AGCTATTATCATCATCACCATCACCACGATTATGATATTCCGACCACCGAAAATCTGTATTTTCAGG
GTGCAATGGTTACCACACTGAGCGGTCTGAGTGGTGAACAGGGTCCGAGCGGTGATATGACCA
CAGAAGAAGATAGCGCAACCCATATCAAATTCAGCAAACGTGATGAAGATGGTCGTGAACTGGC
AGGCGCAACCATGGAACTGCGTGATAGCAGCGGTAAAACCATTAGCACCTGGATTAGTGATGGT
CACGTGAAAGATTTTTATCTGTATCCGGGTAAATATACCTTCGTTGAAACCGCAGCACCGGATGGT
TATGAAGTTGCAACCCCGATTGAATTCACCGTTAACGAAGATGGCCAGGTTACCGTTGATGGTGA
AGCAACCGAAGGTGATGCACATACCGGTAGCAGTGGTAGC

**Forward primer for vector (annealing region highlighted):**

5’- ATACCGGTAGCAGTGGTAGCTAAACTGGCCTCACCGG-3’

Length: 17 nt

GC: 59%

Length: 37

Overhang: 20

**Reverse primer for vector (annealing region highlighted):**

5’- TGGTGATGATGATAATAGCTCATATGTATATCTCCTTCTTAAAGTTAAACA-3’

Length: 31 nt

GC: 26%

Length: 51

Overhang: 20

#### *1.3.2 Expression*

***General.*** Centrifugation was performed with a Sorvall RC-5 centrifuge using SS-34 and GS3 rotors (Watham, MA, USA). Cells were lysed using a Q700 sonicator from Qsonica (Newtown, CT, USA). UV-Vis absorbance and $O_{600}$ measurements were made on a Genesys 150 UV-Vis Spectrometer from Thermo Scientific (Waltham, MA, USA). Protein purification was performed on an ÄKTA Pure 25 Fast Protein Liquid Chromatography (FPLC) (Cytiva, Marlborough, MA) using a Superdex 75 Increase 10/300 column (Cytiva, Marlborough, MA, USA). The dialysis tubing used was made of regenerated cellulose from Spectrum Labs (Waltham, MA, USA). The protein stocks were concentrated using Amicon Ultra-4 10k MWCO Spin Filter and sterile filtered with 0.22 µm PES Millex-GP Syringe Filter both from EMD Millipore (Burlington, MA, USA). Isopropyl β-D-1-thiogalactopyranoside (IPTG) was purchased from LabScientific Inc. (Highlands, NJ, USA). Protease inhibitor cocktail cOmplete mini EDTA-Free tablets were purchased from Roche (Basel, Switzerland). Nickel agarose resin (high density) was purchased from GoldBio (St. Louis, MO, USA). Gels were run utilizing a SDS-PAGE apparatus from Bio-Rad (Hercules, CA, USA). The Spectra Multicolor Low Range Protein Ladder and GelCode Blue Stain Reagent was purchased from Thermo Scientific (Waltham, MA, USA). Gels were imaged using the G:Box Mini from Syngene (Frederick, MD, USA). NuPAGE™ LDS Sample Buffer (4X) was purchased using from Invitrogen/ Thermo Fisher (Waltham, MA, USA).

The pTXB1 $His_6$-SpyCatcher003 plasmid was transformed into BL21(DE3) *E. coli* cells by heat-shock and grown on LB-agar plates with ampicillin (100 µL/mL). Plates were incubated at 37 °C for at least 16 hours. Individual colonies were isolated and grown in 5 mL sterile LB media with ampicillin (100 µL/mL) and grown overnight at 37 °C with 250 RPM shaking. To each liter of sterile LB media up to three primary cultures were added and grown until saturation at 37 °C with 250 RPM shaking. Once an $OD_{600}$ of 0.6-0.8 was reached, the cells were induced with IPTG to a final concentration of 1 mM. The cells were incubated at 18 °C overnight with 250 RPM shaking. The cells were harvested the following day by centrifugation at 4,000 rpm (2,704 *x g*) in a GS3 rotor with a Sorvall RC-5 centrifuge for 20 minutes at 4°C. The supernatant was discarded, and the pellet was resuspended in 30 mL lysis buffer (50 mM Tris-HCl, 300 mM NaCl pH 7.8) with 1 mM PMSF and a broad-spectrum protease inhibitor tablet. The resuspended cells were lysed via sonication (35 amp power, 1 second pulse, 2 second rest for 5 minutes total) and then pelleted at 14,000 rpm (23,426 *x g*) with a SS-34 rotor in a Sorvall RC-5 centrifuge.

#### *1.3.3 Purification*

Nickel agarose resin was added to a fritted column for a settled volume of 5 mL. The resin was equilibrated with 25 mL of lysis buffer. The supernatant of the cell lysate was incubated with the equilibrated resin for at least 60 minutes with rotation at 4 °C. The lysate was applied back to the fritted column and the flow-through was allowed to drain. The resin was washed with 25 mL lysis buffer and 25 mL of lysis buffer with 20 mM imidazole. The desired construct was eluted from the resin with 12 mL of lysis buffer with 300 mM imidazole. DTT was added to a final concentration of 1 mM and 1-2 aliquots of Tobacco Etch Virus (TEV) protease were added to cleave the His-tag by rotation overnight at 4 °C. The sample was transferred to 3.5 kDa MWCO dialysis tubing and was dialyzed into 1x PBS pH 7.5. The non-His-tagged material was isolated with a second-nickel purification. Nickel agarose resin was added to a fritted column for a settled volume of 5 mL. The resin was equilibrated with 25 mL of the dialysis buffer (1x PBS pH 7.5). The sample was incubated with the equilibrated resin for at least 60 minutes with rotation at 4 °C. The resin was applied back to the fritted column and the flow-through containing the non-His-tagged material was collected. The flow-through was spin concentrated to ~2 mL and then dialyzed into 1x PBS pH 7.5.

The construct was further purified via size-exclusion chromatography on a Superdex 75 Increase on an FPLC with 1.2 CV isocratic gradient using 1x PBS pH 7.5. The desired fractions were identified with MALDI using sinapinic acid (SA) matrix and spin concentrated to ~ 50 or 100 μM, aliquoted and stored at -20°C until further use. The concentration was determined based on the UV absorbance ($\varepsilon_{280nm}$ = 17,420 (His$_6$-SpyCatcher003) or 11,460 (SpyCatcher003) $M^{-1}$ $cm^{-1}$).

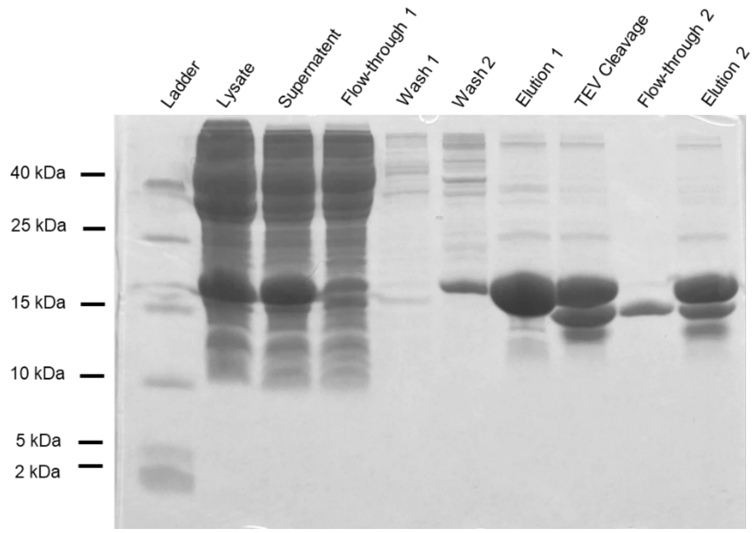


**Figure S2.** 14% Tris-Tricine SDS-PAGE gel of SpyCatcher003 expression and purification.

#### *1.3.4 $^{15}$N-SpyCatcher003*

To generate isotopically labelled material, protein was expressed as described above, but with the following modifications. Instead of LB, minimal media (10x M9 without ammonium chloride = 60 g/L $Na_2HPO_4$, 30 g/L $KH_2PO_4$ and 5 g/L NaCl) and SMM (20 mL = $^{15}NH_4Cl$ (1 g), glucose (2 g), $MgSO_4$ (0.24 g), $CaCl_2$ (dihydrate salt, 15 mg) and yeast extract (200 mg)) media were used. After transformation, a single colony was isolated and added to a 50 mL starter culture containing minimal media (49 mL of 1.5x M9) supplemented with SMM (1 mL), $FeSO_4$ heptahydrate salt (50 µL of 15 mg/mL), $ZnCl_2$ (50 µL of 15 mg/mL) and ampicillin (100 µL/mL final concentration). The starter culture was incubated overnight at 37 °C with 250 RPM shaking. The following morning, 490 mL of autoclaved 1.5x M9 media was supplemented with SMM (10 mL), $FeSO_4$ (0.5 mL of 15 mg/mL), $ZnCl_2$ (0.5 mL of 15 mg/mL) and ampicillin (100 µL/mL final concentration). The starter culture was used to inoculate two 0.5 L of the described media. The rest of the protocol is the same as previously described.

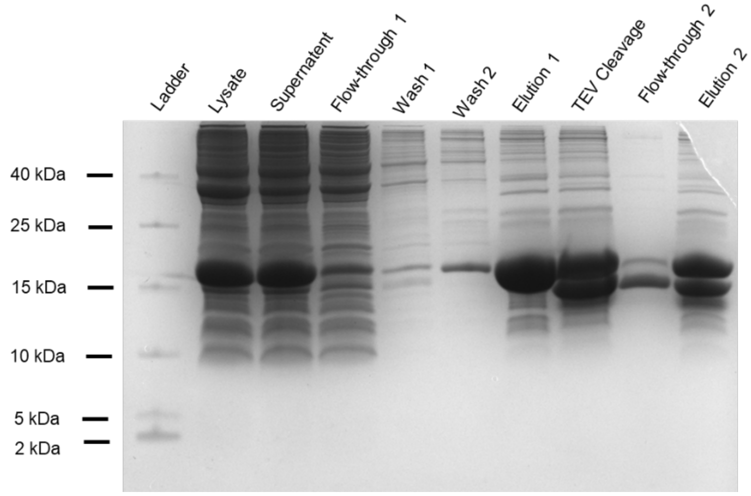


**Figure S3.** 14% Tris-Tricine SDS-PAGE gel of $^{15}$N-SpyCatcher003 expression and purification

**Table S4.** MALDI and characterization of purified SpyCatcher003 and SpyCatcher003/SpyTag003 complexes

| **Construct** | **Calculated MW (Da)** | **Calc. $[M+H]^+$** | **Obs $[M+H]^+$** | **SEC Elution (mL)** |
|---|---|---|---|---|
| SpyCatcher003 | 12,760.44 | 12,761.79 | 12,758.25 | 12.5 |
| $^{15}$N-SpyCatcher003 | 12,904.47 | 12,905.82 | 12,898.70 | 12.4 |
| SpyCatcher003-SpyTag Complex | 14,674.75 | 14,676.10 | 14,676.31 | --- |
| SpyCatcher003-Thio-SpyTag Complexes | 14,690.82 | 14,692.17 | 14,695.42 | --- |
| $^{15}$N-SpyCatcher003-SpyTag Complex | 14,818.78 | 14,820.13 | 14,809.93 | 12.1 |
| $^{15}$N-SpyCatcher003-Thio-SpyTag Complexes | 14,834.85 | 14,836.20 | 14,834.20 | --- |

### 4. Gel Shift Assay

#### *1.4.1 4% Tris-Tricine SDS-PAGE Gel*

The 14% separation gel stack was prepared as outlined and poured after addition of APS and TEMED. The separation layer was topped with water to ensure a level layer. After the separation layer has polymerized, the 4% stacking gel stack was prepared and poured after addition of APS and TEMED. The 1x anode buffer was added inside the gel cassette and the 1x cathode buffer was added outside. Samples were run through the stack layer with 60 V and through the separation layer with 150 V. Gels were prepared and run on the same day.

3x Gel buffer

- 3 M Tris
- Adjust pH to 8.45
- 0.3% sodium dodecyl sulfate (SDS)

10x Cathode buffer

- 1 M Tris

- 1 M Tricine
- 1% SDS

10x Anode buffer

- 2 M Tris
- Adjust pH to 8.9 (with HCl)

14% Separation gel

- 1.75 mL 40% acrylamide
- 1.67 mL 3x gel buffer
- 0.80 mL 50% (v/v) glycerol
- 0.74 mL Milli-Q water
- 50 µL APS

5 µL TEMED

- 4% Stacking gel
- 0.15 mL 40% acrylamide
- 0.50 mL 3x gel buffer
- 0.80 mL Milli-Q water
- 25 µL APS
- 2.5 µL TEMED

#### *1.4.2 Assay*

A 0.05 µmol aliquot (or 0.1 µmol for $Met^{S}_{8}$) of dried SpyTag was dissolved in 1.0 mL of 1x PBS pH 7.5. The concentration was determined based on the UV absorbance ($\varepsilon_{280nm}$ = 2,980 $M^{-1}$ $cm^{-1}$ for oxo SpyTag or $\varepsilon_{274nm}$ = 12,969 $M^{-1}$ $cm^{-1}$ for thioamide-containing SpyTag). This SpyTag sample was further aliquoted and stored at -20°C.

Ligations were performed with 10 µM SpyCatcher and 20 µM SpyTag (2 eq) in 1x PBS pH 7.5 (final volume of 250 µL). After the 14% Tris-Tricine SDS-PAGE Gel was poured, the quench samples were prepared (4x LDS + 5 µL 200 mM BME and 5 µL sterile MilliQ water). Based on the UV determined concentrations, 1x PBS pH 7.5 was added to an Eppendorf tube with a stir-bar to achieve the appropriate final concentrations. SpyCatcher003 from a freshly thawed aliquot was added (10 µM final concentration). To the t = 0 sec timepoint, SpyCatcher003 (0.1 nmol) and SpyTag (0.2 nmol) were added directly to the quench solution. Additionally, enough water was added so that the t = 0 sec quench sample had 20 µL total volume. SpyTag was added (20 µM final concentration) and the timer was started. For each timepoint, 10 µL of the reaction was removed and mixed by pipetting with the quench sample. After the last timepoint, the samples were vortexed and boiled for 10 minutes. Each quench sample (10 µL) was loaded upon the gel against the Spectra Multicolor Low Range Protein Ladder. The gel was washed with MilliQ water and stained with GelCode Blue overnight. The gels were imaged with the GBox:mini gel imager the following day. The reaction was stored at -20 °C.

#### *1.4.3 Quantification*

The gel bands were quantified with Fiji[44]. The percentage of product (SpyCatcher003/Spytag003) formation was determined from the ratio of product band intensity to the total intensity of both the SpyCatcher003 and product bands. Since we anticipated only 10 µM product, we multiplied this ratio by 10 µM to determine the concentration of the product. These data were fitted using GraphPad Prism 7.01 software (San Diego, Ca) to **Equation S1** to determine the second order rate constant (k, $M^{-1}$ $s^{-1}$). $A_0$ is the starting concentration of SpyCatcher003 (M), $B_0$ is the starting concentration of SpyTag003 (M), x is the time (sec), Y is the concentration of product (M), and z is a correction factor.

**Y=(A0*B0*exA0*k*x -exB0*k*x A0*exA0*k*x -B0*exB0*k*x )** **Equation S1**

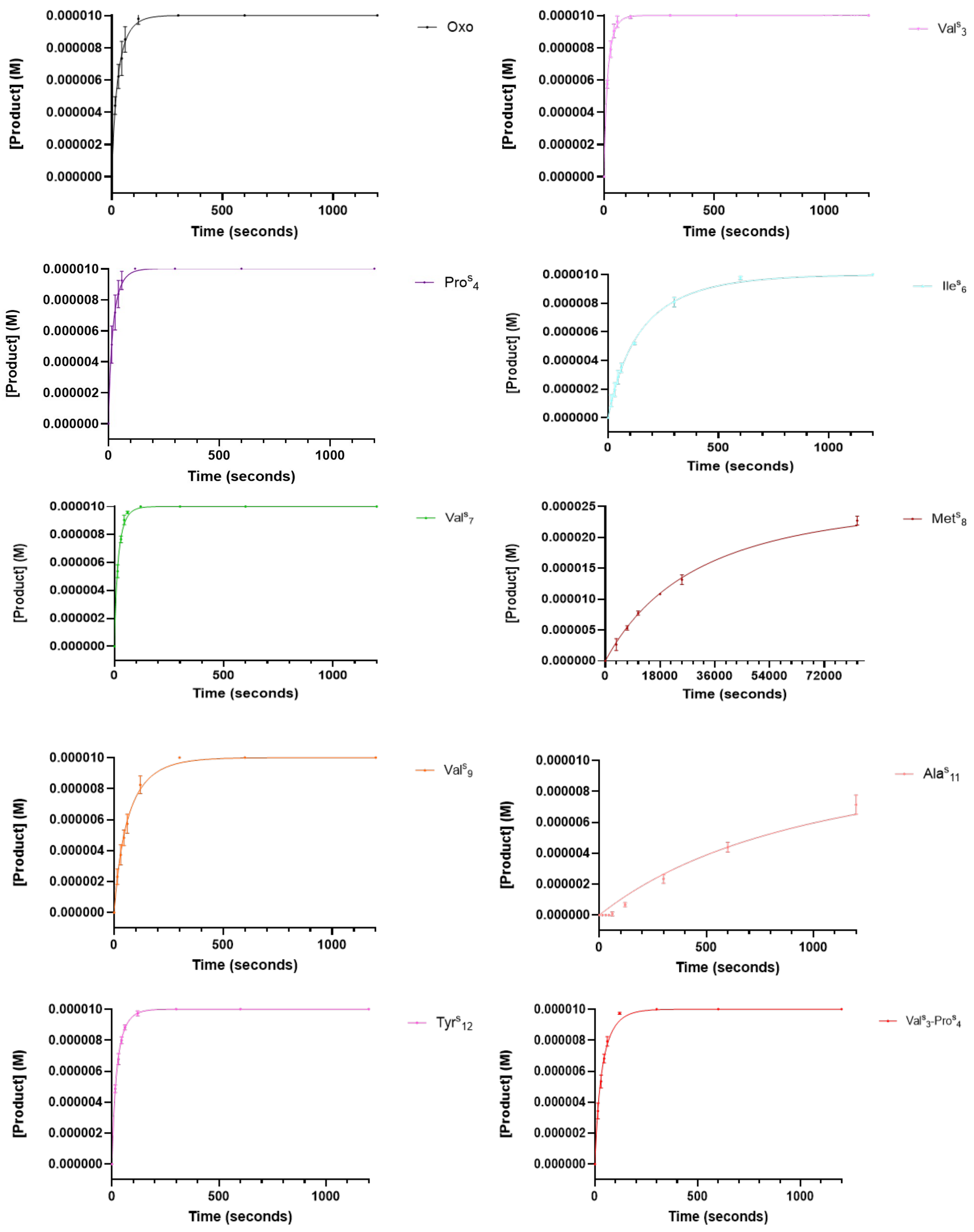


**Figure S4.** Ligation data for Oxo and thioamide SpyTag variants.

### 5. Stopped-Flow Fluorescence Measurements

A 0.05 μmol aliquot of dried SpyTag was dissolved in 1.0 mL of 1x PBS pH 7.5. The concentration was determined based on the UV absorbance ($\varepsilon_{280nm}$ = 2,980 $M^{-1}$ $cm^{-1}$ for oxo SpyTag or $\varepsilon_{274nm}$ = 12,969 $M^{-1}$ $cm^{-1}$ for thioamide-containing SpyTag). SpyCatcher003 aliquot was prepared to be similar in concentration to the SpyTag (~50 μM). The samples and 1x PBS, pH 7.5 were sterile filtered (0.22 μm, PES filter) and stored at -20 °C until use.

Using an AutoSF120 stopped-flow spectrometer from KinTek (Snow Shoe, PA, USA). SpyCatcher003 (25 μM concentration in reaction) and SpyTag (25 μM concentration in reaction) were rapidly mixed and the Trp fluorescence was monitored with excitation at 295 nm. A TECHSPEC fluorescence bandpass filter with a center wavelength of 357.00 nm, a full-width at half-maximum (FWHM) of 44.00 nm, and 75% transmittance from Edmund Optics (Barrington, NJ, USA) was used. 200 μL of sample was added to the well cups and 180 μL was loaded. A shot consisted of 20 μL and three bad shots were discarded once sample was loaded. The PMT HV was set to 500 for all samples and 50 pts/ sec were recorded for each shot. Out of the runs collected, the three replicates with consistent data were averaged and fitted to a one (for $Val^{S}_{3}$) or two-phase decay with GraphPad Prism 7.01 software (San Diego, CA).

Distances from $Trp_{58}$ to the thioamide positions were calculated using PyMol software (New York City, NY). The distance reported is the averaged distance from the carbon of the thioamide bond to δ2 or ε2 carbons of $Trp_{58}$.

### 6. Circular Dichroism

The reactions following the gel shift assays were thawed and excess SpyTag peptide was added to force the reaction to completion. The reaction was stirred at room temperature and complex formation was monitored with MALDI using SA matrix. After completion, the sample was applied to a 3.5 kDa MWCO spin filter, concentrated to ~25 μM and rinsed three times with 1x PBS pH 7.5 to remove unreacted SpyTag peptide. The concentration was determined based on the UV absorbance ($\varepsilon_{280nm}$ = 14,440 $M^{-1}$ $cm^{-1}$ for oxo complex or $\varepsilon_{274nm}$ = 23,946 $M^{-1}$ $cm^{-1}$ for thioamide-containing complex).

The wavelength absorbance scans were collected on a Jasco J-1500 CD spectrometer with a 1 mm path length Helma 110-QS CD cuvette. Measurements were performed at 25 °C, scanning from 350 to 190 nm with a continuous scanning rate of 50 nm/min (1 nm bandwidth and 1 nm data pitch). The instrument was blanked with 1x PBS pH 7.5 prior to sample collection. This blank was manually subtracted from the sample data. The raw signal (θ, mDeg) was converted to the mean residue ellipticity ($\theta_{MRE}$) (**Equation S2**) where l is the pathlength in cm, $n_R$ is the number of residues and c is the concentration in M.

**MRE= c*l*nR** **Equation S2**

The scans were smoothed with GraphPad Prism 7.01 software (San Diego, Ca) by averaging 10 neighbors on each side, and using a fourth order smoothing polynomial.

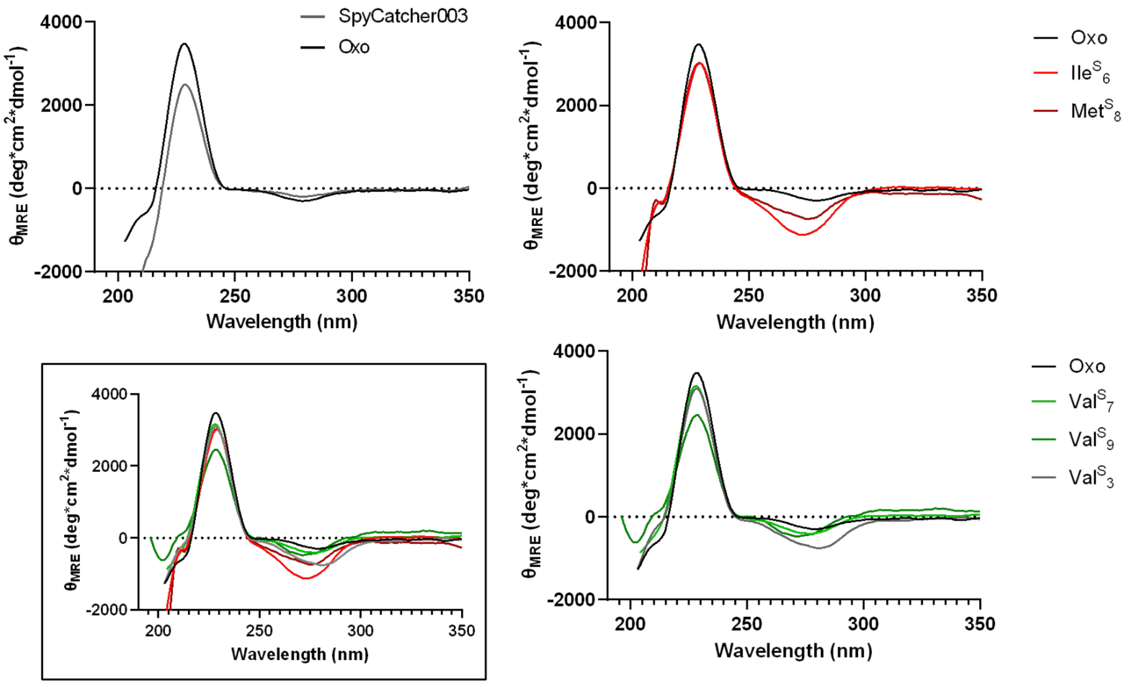


**Figure S5.** CD wavelength scan of SpyCatcher003 alone, Oxo complex, and complexes with internal thioamides (top) and external thioamides (bottom).

Overlay of all spectra is shown in the inset. The positive signature at 230 nm was previously observed for this fold (see Hagan et al.). As previously observed, the thioamide π-π* absorption at 270 nm depends upon the residue and position. Since there is no significant secondary structure as observable by CD, measuring thermostability based on CD is not possible.

### 7. $^1H$-$^{15}N$ HSQC NMR

An aliquot of $^{15}N$-SpyCatcher003 (0.15 µmol) in 1x PBS pH 7.5 was thawed and added to excess dried SpyTag peptide. The reaction was stirred at room temperature and the reaction was monitored with MALDI using SA matrix. When the complex formation was complete, the complex was purified from excess peptide with size-exclusion chromatography on a Superdex 75 Increase column in 1x PBS pH 7.5 for 1.2 CV. The desired fractions were combined and dialyzed into 27.8 mM phosphate pH 6.0 at 4°C with 3.5 kDa MWCO dialysis tubing. Following dialysis, the sample was spin-concentrated with Amicon Ultra-4 10k MWCO Spin Filter to < 1 mL and rinsed with 10 mL of 27.8 mM phosphate pH 6.0. The sample was concentrated to ~110 µM. $D_2O$ was added for a final percentage of 10% (v/v) of the total NMR sample volume. 2-Dimethyl-2-silapentane-5-sulfonate (DSS) was added to each NMR sample as the $^1H$ internal reference. (DSS was added from a 1 mg/mL stock in sterile Milli-Q water to ~50 µM final concentration.) Therefore, the final NMR samples were between 80-100 µM in 25 mM phosphate pH 6.0 containing 10% $D_2O$, 50 µM DSS.

NMR data were collected on a Bruker AVANCE NEO 600 MHz spectrometer. $^1H$-$^{15}N$ HSQC was performed by collecting 2048 points in *f*2, 512 points in *f*1 with 16-32 scans at 298 K. The water $^1H$ signal (ppm) was determined for each sample and inputted as the Transmitted frequency offset (OP1). The spectra were processed with MestReNova 14.1.0 (Santiago de Compostela, Spain). Apodization of Sine Square 90˚ was used for both *f*2 and *f*1, and zero-fill was 2x the size of the FID. A baseline correction of a Berstein polynomial fit of order 3 was used.

### 8. Protein/Peptide Sequences

**$His_6$-SpyCatcher003 (construct post-TEV cleavage)**

MSYYHHHHHHDYDIPTTENLYFQGAMVTTLSGLSGEQGPSGDMTTEEDSATHIKFSKRDEDGREL AGATMELRDSSGKTISTWISDGHVKDFYLYPGKYTFVETAAPDGYEVATPIEFTVNEDGQVTVDGEA TEGDAHTGSSGS [numbering of SpyCatcher003 residues starts at Met post-TEV cleavage site]

**SpyTag**

RGVPHIVMVDAYKRYK

### 9. AlphaFold Structural Modeling

In generating structural models of SpyCatcher-SpyTag complexes, the sequences for SpyCatcher003 and SpyTag003[1] were input as separate sequences into the AlphaFold 3.0 server (https://alphafoldserver.com).[2] The highest confidence prediction was input to PyMOL and compared to the SpyCatcher001-SpytagTa001structure (PDB ID: 4MLI)[3] using the alignment command. Root mean square deviation (RMSD) was determined across all atoms respective to SpyCatcher001-SpytagTa001 and the CnaB2 domain of FbaB, the fibronectin-binding protein of *Streptococcus pyogenes* (PDB ID: 2X5P).[4]

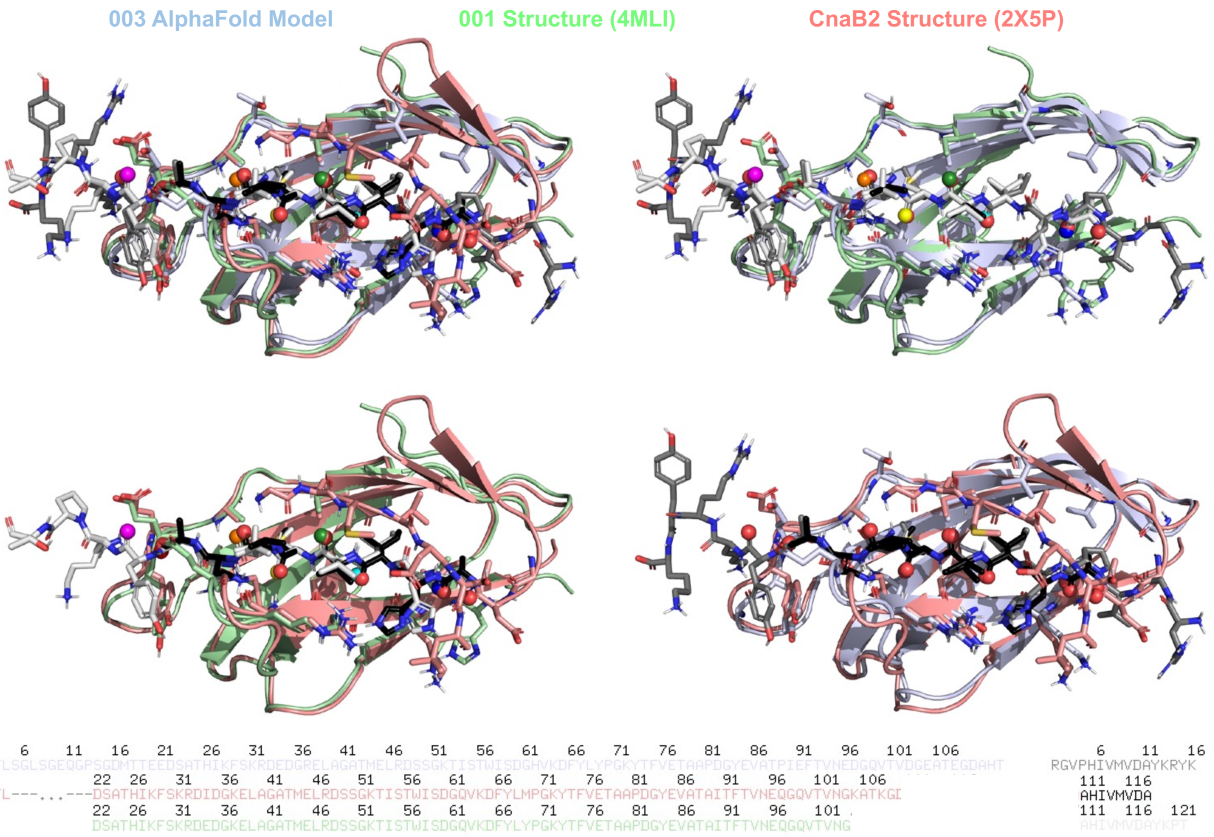


**Figure S6.** Structural models and sequence alignment of SpyCatcher-SpyTag constructs.

**References**

1. Keeble Anthony, H.; Turkki, P.; Stokes, S.; Khairil Anuar Irsyad, N. A.; Rahikainen, R.; Hytönen Vesa, P.; Howarth, M., Approaching infinite affinity through engineering of peptide–protein interaction. *Proc. Natl. Acad. Sci. U. S. A.* **2019,** *116* (52), 26523-26533.

2. Abramson, J.; Adler, J.; Dunger, J.; Evans, R.; Green, T.; Pritzel, A.; Ronneberger, O.; Willmore, L.; Ballard, A. J.; Bambrick, J.; Bodenstein, S. W.; Evans, D. A.; Hung, C.-C.; O'Neill, M.; Reiman, D.; Tunyasuvunakool, K.; Wu, Z.; Žemgulytė, A.; Arvaniti, E.; Beattie, C.; Bertolli, O.; Bridgland, A.; Cherepanov, A.; Congreve, M.; Cowen-Rivers, A. I.; Cowie, A.; Figurnov, M.; Fuchs, F. B.; Gladman, H.; Jain, R.; Khan, Y. A.; Low, C. M. R.; Perlin, K.; Potapenko, A.; Savy, P.; Singh, S.; Stecula, A.; Thillaisundaram, A.; Tong, C.; Yakneen, S.; Zhong, E. D.; Zielinski, M.; Žídek, A.; Bapst, V.; Kohli, P.; Jaderberg, M.; Hassabis, D.; Jumper, J. M., Accurate structure prediction of biomolecular interactions with AlphaFold 3. *Nature* **2024,** *630* (8016), 493-500.

3. Li, L.; Fierer, J. O.; Rapoport, T. A.; Howarth, M., Structural Analysis and Optimization of the Covalent Association between SpyCatcher and a Peptide Tag. *Journal of Molecular Biology* **2014,** *426* (2), 309-317.

4. Oke, M.; Carter, L. G.; Johnson, K. A.; Liu, H.; McMahon, S. A.; Yan, X.; Kerou, M.; Weikart, N. D.; Kadi, N.; Sheikh, M. A.; Schmelz, S.; Dorward, M.; Zawadzki, M.; Cozens, C.; Falconer, H.; Powers, H.; Overton, I. M.; van Niekerk, C. A. J.; Peng, X.; Patel, P.; Garrett, R. A.; Prangishvili, D.; Botting, C. H.; Coote, P. J.; Dryden, D. T. F.; Barton, G. J.; Schwarz-Linek, U.; Challis, G. L.; Taylor, G. L.; White, M. F.; Naismith, J. H., The Scottish Structural Proteomics Facility: targets, methods and outputs. *Journal of Structural and Functional Genomics* **2010,** *11* (2), 167-180.
